# Methotrimeprazine exerts antiviral and neuroprotective effects in Japanese encephalitis virus infection through activation of adaptive ER stress and autophagy

**DOI:** 10.1101/2023.04.05.535645

**Authors:** Surendra K. Prajapat, Laxmi Mishra, Sakshi Khera, Shadrack D. Owusu, Kriti Ahuja, Puja Sharma, Rajan Singh, Dinesh Mahajan, Arup Banerjee, Rajender K. Motiani, Sudhanshu Vrati, Manjula Kalia

## Abstract

Japanese encephalitis virus (JEV) is the leading global cause of virus-induced encephalitis. Its pathogenesis is driven by a combination of neuronal cell death and neuroinflammation. We hypothesized that pharmacological upregulation of autophagy could exert a neuroprotective antiviral effect, and tested a panel of forty-two FDA-approved drugs that were shown to induce autophagy. Four drugs were tested in the JE mouse model based on in vitro protective effects on neuronal cell death, inhibition of viral replication, and anti-inflammatory effects in microglial cells. The antipsychotic phenothiazines Methotrimeprazine (MTP) and Trifluoperazine (TFP) showed a significant survival benefit with reduced virus titers in the brain, prevention of blood-brain barrier (BBB) breach, and inhibition of neuroinflammation. Both drugs were potent mTOR-independent autophagy flux inducers. Mechanistically MTP inhibited SERCA channel functioning, thereby resulting in rise in cytosolic calcium levels, and induction of a unique adaptive ER stress response. In virus infected drug treated cells, there was a strong transcriptional downregulation of type I interferon and interferon-stimulated genes and upregulation of cholesterol metabolic pathway genes. The drugs exerted an autophagy-dependent antiviral effect at the level of JEV protein translation/replication complex formation in diverse cell types. Inhibition of inflammatory cytokine/chemokine release from mouse microglial cells was partly autophagy-dependent. Our study suggests that MTP exerts a combined antiviral and anti-inflammatory effect in JEV infection, and has therapeutic potential to be repurposed for JE treatment.

## Introduction

Japanese encephalitis virus (JEV) belongs to the Flaviviridae family that includes several pathogenic arboviruses such as West Nile virus (WNV), Yellow fever virus and Dengue virus (DENV). JEV is transmitted by infected Culex mosquitoes and is maintained through an enzootic cycle between birds, pigs and other vertebrate hosts. The disease is endemic in south-east Asian countries including India, with both epidemic and sporadic occurrences [1]. Over the years the virus has shown significant geographical expansion into regions not previously reported to have JE [2].

JEV is neurotropic, and its clinical manifestations range from mild febrile illness to encephalitis and death [3]. The pediatric population is most severely affected, and treatment is mostly supportive with no effective antiviral therapy available [4]. Virus induced perivascular and central nervous system (CNS) inflammation is linked to elevated intracranial pressure, seizures, movement disorders and flaccid paralysis [1, 5, 6]. Neuronal damage to the thalamus and brain stem often results in permanent neurological sequelae among the survivors [7].

Following inoculation through a mosquito bite, the virus first replicates in the local dermal cells such as fibroblasts, endothelial cells and tissue-resident dendritic cells (DCs), and spreads to local lymph nodes and other organs [1, 8]. The virus also replicates in monocytes/macrophages and in most cases, is cleared by an effective peripheral immune response [9–11]. The virus can invade the CNS either through basolateral release from infected brain microvascular endothelial cells (BMECs), or diapedesis of infected peripheral immune cells. JEV replicates efficiently in neurons, microglia and astrocytes. Production of inflammatory cytokines and metalloproteases by JEV infected BMECs, microglia, and astrocytes triggers the degradation of tight junction proteins leading to loss of brain endothelial barrier permeability. Studies have shown that blood-brain barrier (BBB) breach is not a cause, but a consequence of virus infection of the CNS and neuroinflammation [12]. Neuronal cell death, which is augmented by neuroinflammation is the major driver of pathogenesis [13].

JEV being an RNA virus, replicates in close association with ER derived membranes, and results in the activation of stress responses such as the unfolded protein response (UPR), ER stress, generation of ROS, and upregulation of autophagy [14–18]. In the context of JEV, cellular autophagy is upregulated through the activation of ER and oxidative stress, and functions primarily as an antiviral mechanism by restricting virus replication and cell death [16–18]. At later time points of infection, autophagy dysregulation is observed, which enhances virus induced neuronal death. This lead us to hypothesize that autophagy upregulation could inhibit virus replication, neuronal cell death and neuroinflammation, and is thus likely to be neuroprotective.

Established defects in autophagy in conditions such as cancer, neurodegeneration, inflammation and metabolic disorders have lead researchers to focus on the discovery of novel drugs/compounds that can modulate autophagy. Autophagy upregulation has also been shown to have therapeutic potential for neurodegenerative diseases [19, 20]. Several FDA-approved drugs have been shown to enhance autophagy. Therefore, they have the potential to be repurposed in disease conditions where autophagy upregulation is likely to be beneficial.

Here we have examined FDA-approved drugs with autophagy inducing potential for their effect on JEV infection in vitro and in mouse model of disease. The typical antipsychotic drugs of the phenothiazine family Methotrimeprazine (MTP) and Trifluoperazine (TFP) showed strong inhibition of virus replication, and microglial inflammation, along with significant protection in the mouse model of disease. These drugs induced mTOR-independent functional autophagy flux, and their antiviral activity was observed to be autophagy dependent. MTP treatment resulted in calcium dysregulation, eIF2α phosphorylation and a unique adaptive ER stress gene signature. There was a strong downmodulation of type I interferon (IFN) and upregulation of cholesterol metabolic genes. The inhibition of proinflammatory cytokine secretion from microglial cells was partly Atg5 dependent. Our study suggests that MTP induces autophagy and adaptive ER stress in diverse cell types creating an antiviral and neuroprotective environment during JEV infection.

## Results

### Primary screening of autophagy inducing FDA approved drugs for antiviral and anti-inflammatory effects in vitro

Studies from our laboratory have shown that JEV infection induced cellular stress responses such as ER and oxidative stress, result in the activation of the UPR and autophagy, that play crucial roles in regulating JEV replication and cell death [16–18]. Since autophagy upregulation has potential neuroprotective roles, we tested a panel of FDA-approved drugs that have been documented as autophagy-inducers, for any anti-JEV effect. The study was initiated with forty-two drugs (Table S1), and all were tested at a concentration of 10 µM which was established as non-cytotoxic in the mouse Neuro2a cell line (Fig S1A). A primary screening was performed using virus induced neuronal cell death assay. JEV infection of Neuro2a cells results in MOI and time-dependent cell death. A 5 MOI infection for 48 h, which results in ∼ 80% cell death was chosen for the assay (Fig S1B). From the panel ten drugs: Bromhexine, Clonidine, Flubendazole, Fluoxetine, Lithium chloride, Memantine, Metformin, MTP, Rilmenidine and Sodium valproate showed reduction in JEV-induced cell death (Fig S1C), and were shortlisted for further studies. All drugs resulted in a significant reduction in JEV RNA levels (Fig 1A), and four drugs: Clonidine, Fluoxetine, Metformin and MTP significantly reduced virus titers in Neuro2a cells (Fig 1B).

**Fig 1:**
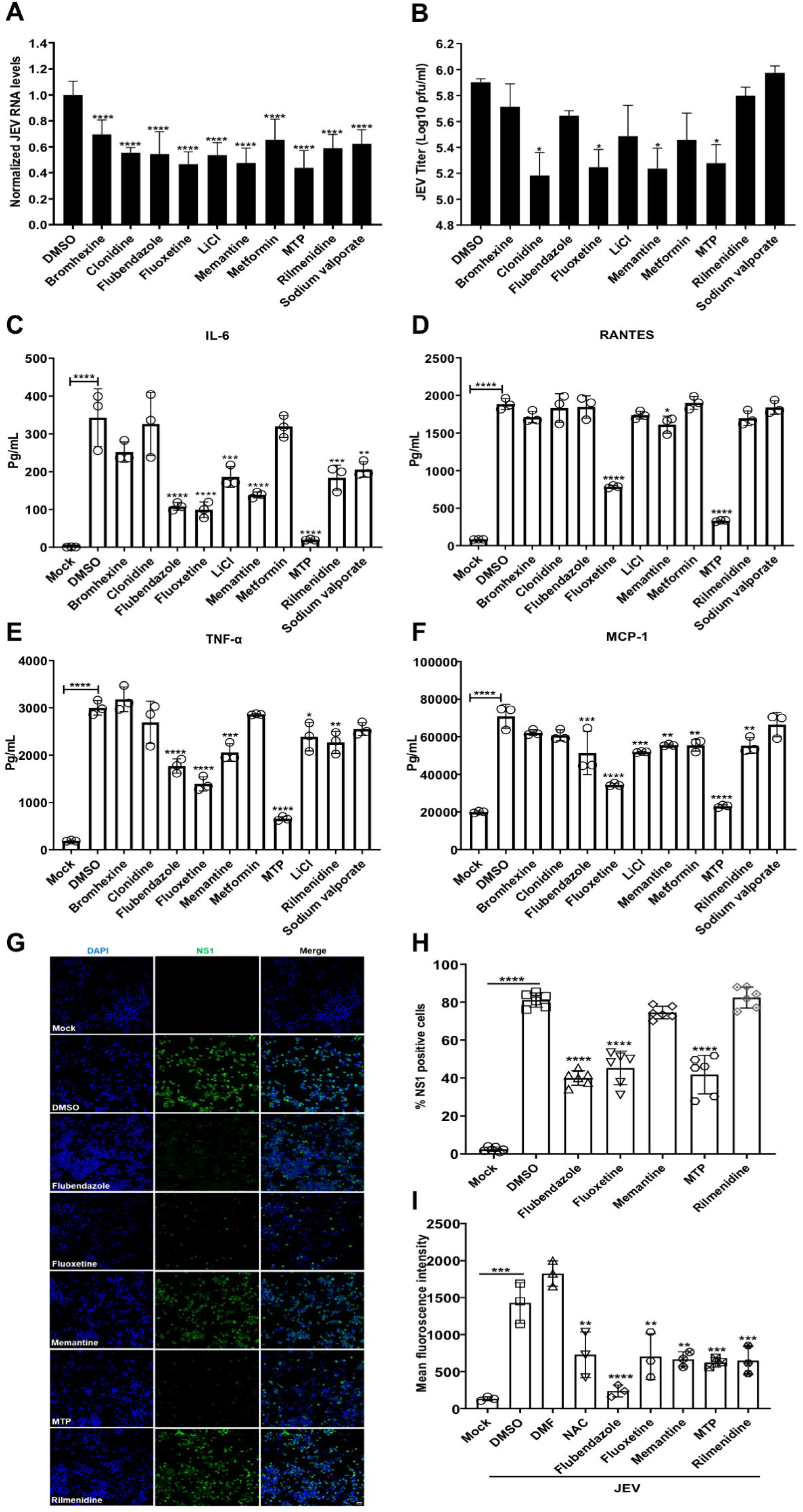
Antiviral and anti-inflammatory effect of FDA-approved drugs against JEV infection. (A-B) Neuro2a cells were infected with JEV (MOI 1), and at 1 hpi treated with DMSO/drugs (10 µM). (A) Cells were harvested at 24 hpi and viral RNA levels were quantified using qRT-PCR. Graph shows the relative expression levels of JEV RNA normalized to DMSO-treated control. (B) Culture supernatant was collected to determine extracellular virus titer using plaque assays. Data represents values obtained from three independent experiments. (C-F) N9 cells were mock/JEV-infected (MOI 1), and at 12 hpi treated with DMSO/ drugs (10 µM) for 24 h. Cytokine concentrations (pg/ml) were quantified from the culture supernatant using CBA assay. Data shows values from one representative experiment with biological triplicates. (G-H) Neuro2a cells were mock/JEV (MOI 5) infected, and at 1 hpi were treated with DMSO/ drugs (10 µM) till 24 hpi. (G) Cells were immunostained for JEV NS1 (green) and images were acquired on high-content imaging system. Scale bar, 10 µm. (H) Bar-graph showing percentage of NS1 positive cells. (I) Neuro2a cells were mock/JEV (MOI 5) infected, and at 1 hpi were treated with DMSO/ FDA-drugs (10 µM); or at 16 hpi treated with DMF (70 µM)/ NAC (3 mM), and maintained till 24 hpi. Post-treatment, cells were stained with oxidative stress indicator CM-H2DCFDA and fluorescence intensity was measured using flow cytometry. The graph represents mean fluorescence intensity values. All data are expressed as means ± SD, statistical significance was determined using one-way ANOVA. *, P<0.05; **, P<0.01; ***, P<0.001; ****, P<0.0001.

As previously reported [13], we observed that JEV infects microglial cells efficiently (Fig S2A) and results in robust secretion of pro-inflammatory cytokines such as IL-6, RANTES, TNF-α and MCP-1 (Fig S2B-E). After establishing non-cytotoxic concentrations (Fig S2F), the ten short-listed drugs were checked for their effect on JEV replication (Fig S2G), and inhibition of pro-inflammatory cytokine release from infected N9 microglial cells (Fig 1C-F).

Based on observations from both neuronal and microglial cells, five drugs appeared as promising antivirals: Flubendazole, Fluoxetine, Memantine, MTP and Rilmenidine. These drugs were further tested for their effect on JEV protein translation/replication complex formation (Fig 1G-H), and ROS production (Fig 1I) in Neuro2a cells and a significant inhibition was observed. MTP treatment also showed a significant inhibition of cytokine release from LPS treated microglial cells (Fig S2H).

### Phenothiazines exert an antiviral and anti-inflammatory effect and show protection in JEV mouse model

We next tested Flubendazole, Fluoxetine, MTP and Rilmenidine in a JEV infected C57BL/6 mouse survival assay (Fig S3). A mouse adapted isolate JEV-S3 was used, which results in development of typical encephalitis symptoms: loss of weight, body stiffening, piloerection, hind limb paralysis by 5-6 days post-infection (dpi), and death within 2-3 days of symptom onset [21]. The JEV infected mice developed typical disease symptoms and showed a median survival time (MST) of 8-9 days (Fig S3, 2A-B). Brain viremia was detectable by 3 dpi and increased rapidly thereafter till 6 dpi indicative of active infection (Fig 2C). The Evans Blue (EB) leakage test showed that the BBB was intact at 3 dpi, clearly indicating that the breach is not required for virus neuroinvasion. The infected mice showed barrier permeability by 6 dpi (Fig 2D-E). Strikingly, the MTP treated mice showed a significant survival benefit (Fig S3, 2A-B), with delayed virus invasion into the brain, and significantly lower viremia (Fig 2C). The drug treated mice also showed complete protection of the barrier that was comparable to control uninfected mice (Fig 2D-E). These data demonstrate that MTP exerted a significant neuroprotective effect in the JE mouse model.

**Fig2:**
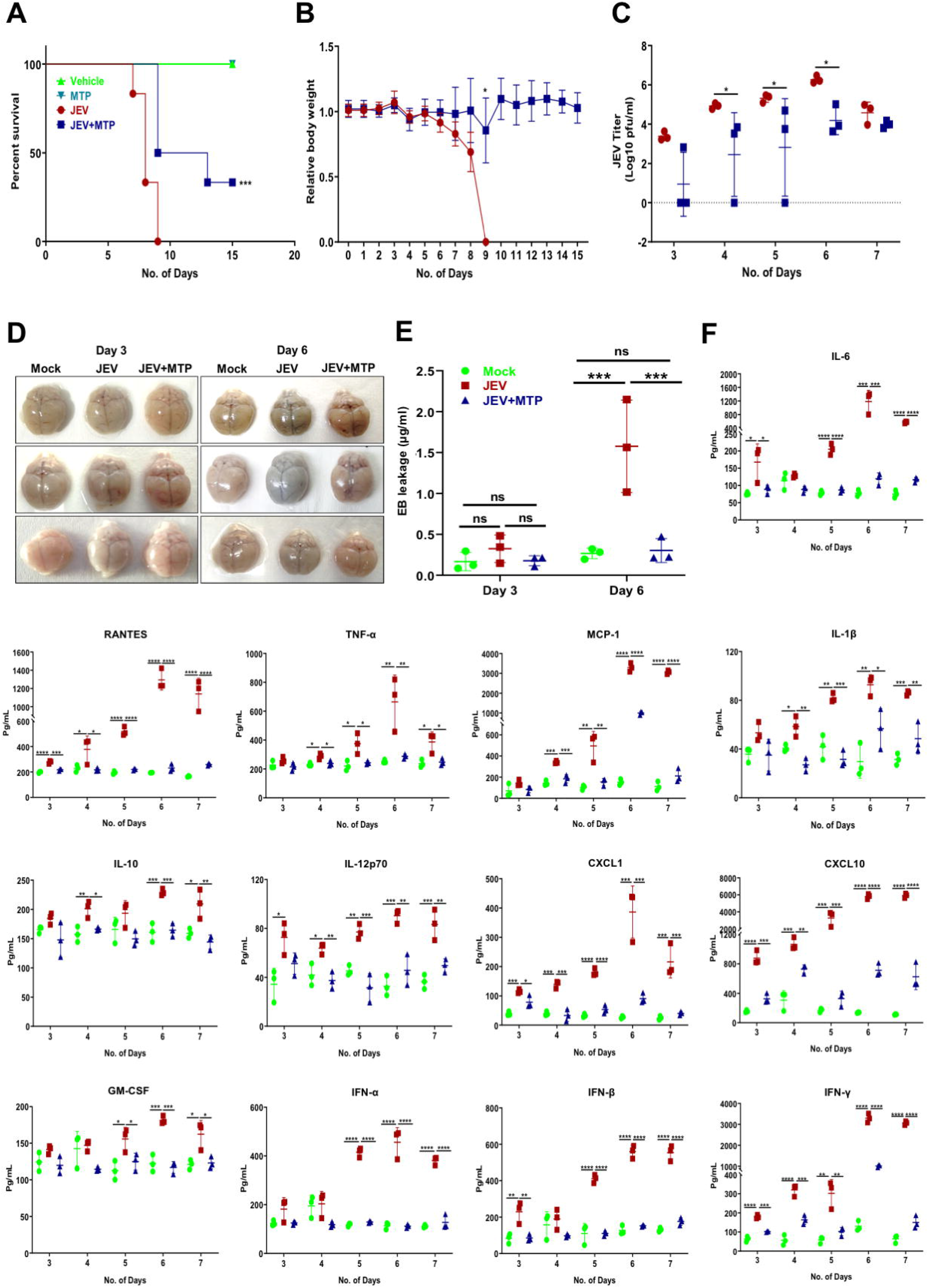
Efficacy of MTP in JEV-mouse model. 3 week old C57BL/6 mice were mock/ JEV-S3 (10^7^ pfu) infected through an i.p. injection, and at 4 hpi were treated with vehicle control (PEG400) or MTP (2mg/kg) by oral gavage at an interval of 24 h for 15 days. All mice were monitored for the appearance of encephalitis symptoms until death. (A) Survival curve of mock (n=4)/MTP (n=4)/JEV (n=6)/JEV+MTP (n=6) was plotted, a Log-rank (Mantel-Cox) test was used to determine the statistical significance comparing JEV and JEV+MTP mice group. (B) Graph representing the change in body weight of vehicle/MTP-treated infected mice group normalized to mock-infected mice group, compared by paired student t-test. (C) Mock or Vehicle/MTP treated infected mice (n=3 /time point from each group) were sacrificed at indicated time points. Brain tissues were homogenized and the supernatant was used for JEV titration using plaque assay. Each data point denotes one mouse, and the virus titer between JEV and JEV+MTP group was compared by paired student t-test. (D-E) Mock or Vehicle/MTP treated infected mice received an i.p. injection of 2% Evans blue dye (100 µl). Mice were sacrificed at 3 and 6 dpi. (D) Representative images showing Evans blue dye distribution in the brain (n=3). (E) The brain tissues were homogenized in DMF (200 mg/50011μl DMF) and absorbance was measured at 62011nm. The concentration of Evans blue was quantitated according to standard curve and significance was compared by two-way ANOVA test. (F) Brain tissue was collected at indicated time points, and an equal amount of protein (30 µg) from each sample was used for the quantitation of cytokine levels using CBA. Data were analyzed with LEGENDplex^TM^ Multiplex assay software, and significance was compared by one-way ANOVA test. All data expressed as means ± SD. *, P<0.05; **, P<0.01; ***, P<0.001;****, P<0.0001.

Since the BBB breach is linked to virus induced neuroinflammation, we tested the levels of several cytokines, chemokines, and interferons in the brains of infected and drug treated mice (Fig 2F). JEV infected mice showed very high levels of proinflammatory cytokines and chemokines: IL-6, TNF-α, RANTES, MCP-1, CXCL-1, CXCL-10, GM-CSF, IL1-β and IFN-β starting at 3 dpi and these increased rapidly peaking at 6-7 dpi (Fig 2F). The anti-inflammatory cytokines IL-10 and IL-12p70 along with IFN-γ were also secreted at high levels in the infected mice, indicative of active T cell infiltration in the brain during infection. Importantly, these effects were completely ameliorated in the drug treated mice indicative of a significant protection from neuroinflammation (Fig 2F).

Reduced neuroinvasion in drug treated mice suggested that some protection was also conferred at the periphery. We tested different doses of MTP for toxicity on bone-marrow derived macrophages (BMDMs) (Fig 3A). Drug treatment of JEV-infected BMDMs resulted in a significant reduction of both virus replication (Fig 3B) and production of several inflammatory cytokines and interferons (Fig 3C). This reduction cannot be attributed entirely to reduced virus replication as a similar inhibition of pro-inflammatory cytokine release was also observed from LPS treated BMDMs (Fig S4), indicating a strong anti-inflammatory effect of MTP.

**Fig3:**
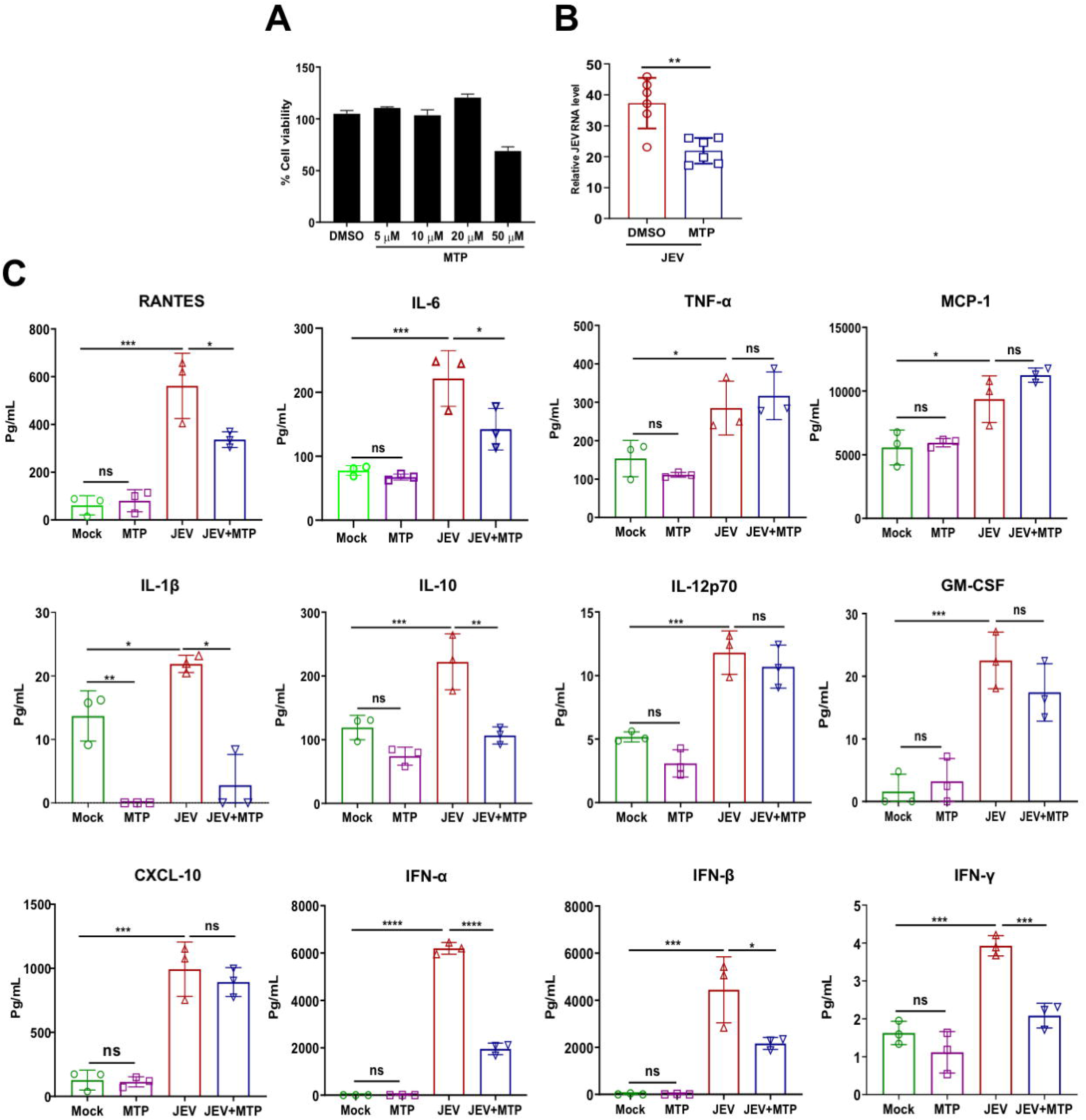
MTP inhibits the secretion of proinflammatory cytokines from JEV infected BMDMs. (A) BMDMs were treated with DMSO/MTP at indicated concentration for 24 h, MTT assay was used to calculate % cell viability. (B-C) BMDMs were mock/JEV (MOI 2) infected for 1 h, followed by DMSO/MTP (10 µM) treatment till 24 hpi. (B) Viral transcript levels from were measured using qRT-PCR and normalized to DMSO-treated infected control, (n=6). (C) Culture supernatant was used for the quantitation of cytokine levels by flow cytometry-based CBA assay (n=3). Similar trends were seen in two independent experiments. All data were expressed as means ± SD and one-way ANOVA test was used for the determination of statistical significance, *, P<0.05; **, P<0.01; ***, P<0.001; **** P < 0.0001, ns; non-significant.

MTP is a widely used FDA-approved antipsychotic that belongs to the phenothiazine class of drugs (Fig 4A). MTP showed an IC50 in the range of 3-3.4 μM in mouse neuronal cells and primary cortical neurons, indicating a strong antiviral response at low doses (Fig 4B-C). Encouraged by our observations, we tested another FDA-approved and widely used phenothiazine drug-Trifluoperazine (TFP) (Fig 4A). This drug also showed robust inhibition of JEV replication with an IC50 of 2 μM (Fig 4D), and a very significant block in viral protein translation/replication complex formation (Fig 4E-F), and production of infectious virus particles (Fig 4G). This drug also exerted a potent anti-inflammatory effect and significantly blocked the release of inflammatory cytokines from virus infected microglial cells (Fig 4H). The drug was also tested in the JE mouse model using a sublethal dose of JEV, and similar to MTP a significant survival benefit was observed (Fig 4I). Collectively these data indicate that phenothiazines exert strong antiviral and anti-inflammatory effect for JEV infection both in vitro and in vivo.

**Fig4:**
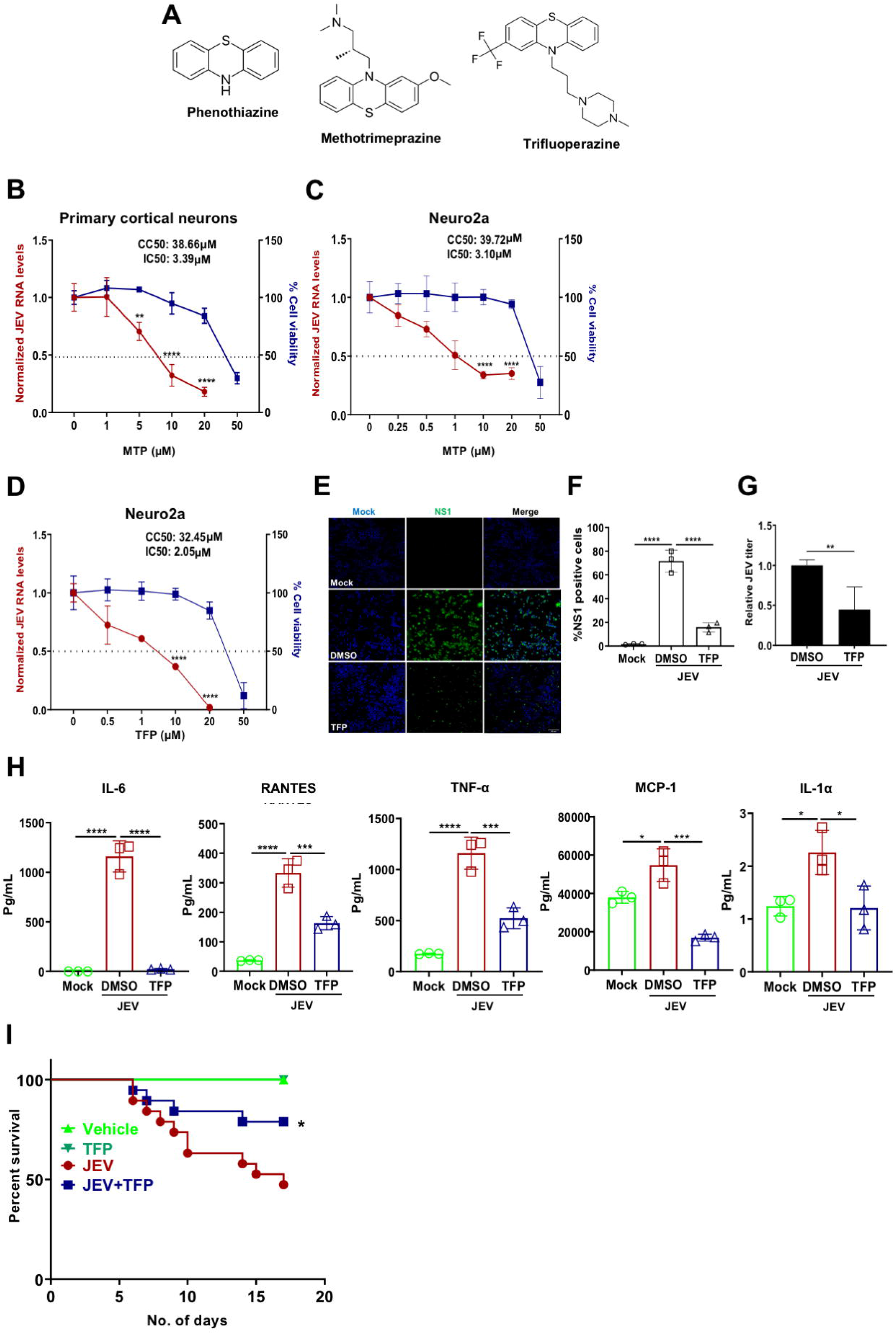
Phenothiazines exert antiviral and anti-inflammatory effects. (A) Chemical structure of the phenothiazine ring and its derivates Methotrimeprazine and Trifluoperazine. (B-C) Primary cortical neurons (B), and Neuro2a cells (C), were mock/JEV (MOI 1) infected, and at 1 hpi were treated with DMSO/MTP at indicated concentrations till 24 hpi. Cells were harvested, qRT-PCR was performed for the quantitation of JEV RNA levels, and % cell viability was measured using MTT assay. Data were normalized to DMSO control, and compared between DMSO and MTP treated JEV-infected cells using unpaired student t-test. Graphs represent the CC50 and IC50 values. (D) Neuro2a cells were mock/JEV (MOI 1) infected and treated with DMSO/TFP at the indicated concentrations till 24 hpi. CC50 and IC50 were calculated as described for panel C. (E-F) Neuro2a cells were mock/JEV (MOI 5) infected, and at 1 hpi treated with 10 µM TFP till 24 hpi. Cells were immunostained with JEV NS1 antibody, and images were visualized by high-content imaging system, representative images are shown. Scale bar, 10 µm (E). (F) Graph shows % NS1 positive cells. (n=3), one-way ANOVA test. (G) Supernatants of DMSO/TFP-treated JEV infected cells was used for the determination of JEV titer using plaque assay. Data is representative of two independent experiments and were compared between DMSO and TFP using paired student t-test. (H) N9 cells were infected with JEV (MOI 1), at 12 hpi cells were treated with TFP (10 µM) for 24 h. Cytokines were quantified from the soup using CBA. (n=3), one-way ANOVA test. (I) C57BL/6 (3 week old) mice were infected by i.p. injection of DMEM (mock) or JEV (10^6^ pfu), at 4 hpi, treated with vehicle control (PEG400)/TFP (1 mg/kg) by oral route at an interval of 24 h till 15 days. All mice were monitored for JEV symptoms until death. The survival curve of mock (n=4)/TFP (n=4)/JEV (n=19)/JEV+TFP (n=19) was plotted from two independent experiments. Log-rank (Mantel-Cox) test was used to determine the statistical significance comparing vehicle and drug-treated infected mice group. All data expressed as means ± SD. *, P<0.05; **, P<0.01; ***, P<0.001; ****, P<0.0001.

### Phenothiazines are mTOR independent autophagy inducers

There is evidence in literature that phenothiazines are autophagy inducers [22, 23]. We also assessed autophagy induction by MTP and TFP in our experimental setup, and observed that these drugs lead to rapid accumulation of lipidated LC3 in Neuro2a cells (Fig 5A-B, S5A-B), and primary cortical neurons (Fig 5C-D), at levels comparable to the autophagy inducer Torin1. Bafilomycin (Baf) A1 treatment in drug treated cells led to further increase of LC3-II levels indicative of functional autophagy flux (Fig 5E-F). This was also confirmed using GFP-LC3-RFP-ΔG expressing reporter MEFs which enable high throughput measurement of GFP/RFP ratio as an indicator of autophagy flux (Fig S5C-D). While high autophagy flux results in low GFP/RFP ratio as seen with Torin1 treatment, a block in flux results in a higher ratio as seen with BafA1. The GFP/RFP ratios indicated that both the drugs were inducing high autophagy flux (Fig S5C-D). Levels of p62 also showed a reduction similar to Torin1 (Fig 5G, H). LysoTrackerRed staining distribution and intensity in drug treated cells was also similar to Torin1 treatment (Fig S5E-F), and the LysoSensor Yellow-Blue assay showed no change in lysosome acidification (Fig S5G-H).

**Fig5:**
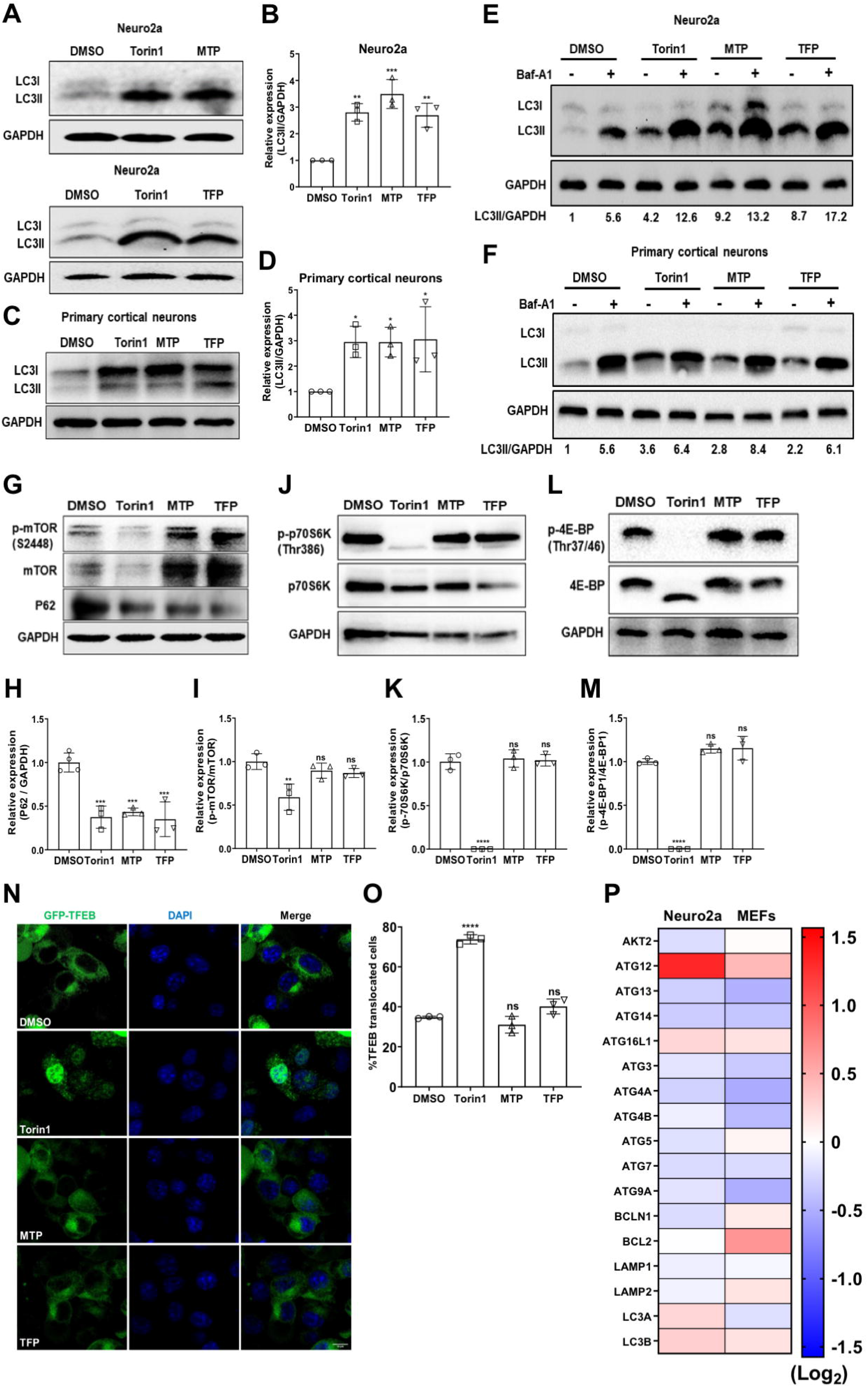
Phenothiazines are mTOR independent autophagy inducers. (A-D) Neuro2a (A-B), and primary cortical neurons (C-D), were treated with DMSO/Torin1 (1 µM)/MTP (10 µM) /TFP (10 µM) for 6 h. Protein lysates were immunoblotted for LC3 and GAPDH (loading control). (B & D) Bar-graph shows relative expression of LC3II/GAPDH normalized to DMSO control from three independent experiments. (E-F) Neuro2a (E), and primary cortical neurons (F), were treated with DMSO/Torin1 (1 µM)/MTP (10 µM) /TFP (10 µM) for 4 h, followed by BafA1 (100 nM) treatment for 2 h. The values below the blot show relative levels of LC3II/GAPDH protein after normalization to DMSO-treated cells. (G-M) Neuro2a cells were treated with DMSO/Torin1 (1 µM)/MTP (10 µM) /TFP (10 µM) for 6 h, protein lysates were analyzed by western blotting with P62 (G, H), p-mTOR (S2448), mTOR (G, I) p-P70S6K (Thr386), p70S6K (J-K), p-4E-BP (Thr37/46), 4E-BP (L-M) and GAPDH (loading control) antibodies. (H, I, K, M) Bar-graphs show relative protein expression level calculated after normalization to DMSO control from three independent experiments. (N-O) EGFP-TFEB expressing Neuro2a cells were treated with DMSO/Torin1 (1 µM)/MTP (10 µM) /TFP (10 µM) for 6 h. Images were acquired on high content imaging system and representative images are shown. Scale bar, 10 µm. (O) Bar-graph showing the percentage EGFP-TFEB nuclear translocation, (n=3). (P) Neuro2a/MEFs were treated with MTP (10 µM) for 6 h and mRNA levels of autophagy genes were determined by qRT-PCR. The heatmap shows relative gene expression level after normalization to DMSO treated control. All data were expressed as means ± SD, one-way ANOVA test was used to calculate statistical significance *, P<0.05; **, P<0.01; ***, P<0.001; ****, P<0.0001; ns, not significant.

A few studies have suggested that another widely used phenothiazine, chlorpromazine (CPZ) inhibits Akt/mTOR [24], and stimulates TFEB nuclear translocation and expression of autophagy-lysosomal target genes [25]. However, we did not observe any mTOR inactivation, as the phosphorylation of mTOR (Fig 5G, I), and its downstream targets p70S6Kinase (Fig 5J, K) and 4EBP (Fig 5L, M), remained unaffected. These drugs also did not lead to any significant TFEB nuclear translocation as was observed with Torin1 treatment (Fig 5N-O). These data suggested that MTP and TFP induce autophagy through an mTOR independent mechanism. We also checked if MTP induced any changes in the transcript levels of autophagy genes, and observed enhanced levels of Atg12, Atg16L1, LC3A, & LC3B in neuronal cells. MEFs also showed a similar transcriptional upregulation of these genes, along with Bcl2 (Fig 5P). Interestingly, both cell types showed downregulation of Atg13, Atg14, Atg3, Atg4a, Atg4b & Atg9 (Fig 5P). Transcriptional upregulation of Atg12 and LC3 has been reported to specifically occur through activated PERK/ATF4, hinting that autophagy activation by MTP could be a result of ER stress activation [26, 27].

### Phenothiazines induce adaptive ER stress and dysregulation of intracellular calcium signaling

MTP treatment resulted in transcriptional changes of autophagy genes that were reminiscent of the PERK/ATF4 pathway activation in response to ER stress [26]. We thus tested other parameters of the UPR in drug treated neuronal cells and MEFs. A primary response is the PERK mediated phosphorylation of eIF2α, that inhibits ribosome ternary complex recycling and attenuates protein translation. Thapsigargin-treated Neuro2a cells showed very rapid and robust eIF2α phosphorylation starting at 1 h of treatment, which was sustained at 3 h, but declined thereafter and returned to baseline by 12 h (Fig 6A). These cells also showed PERK phosphorylation (indicated by mobility shift of PERK) till 6 h (Fig 6A). PERK activation also results in enhanced ATF4 levels that activate a transcriptional response program that can either be adaptive (through autophagy induction) or apoptotic (through production of CHOP and activation of pro-apoptotic proteins) [28, 29]. The other two ER stress sensors are IRE1α that activates XBP1 through an unconventional splicing (XBP1 spl), and ATF6 that is cleaved in the Golgi to generate the ATF6 (N) transcriptional factor. In Thapsigargin treated Neuro2a cells we observed significant transcriptional activation of ATF4, CHOP, GRP78, XBP1, XBP1(s) & ATF6 (Fig 6B), and CHOP protein levels were detectable by 12 h (Fig 6A).

**Fig6:**
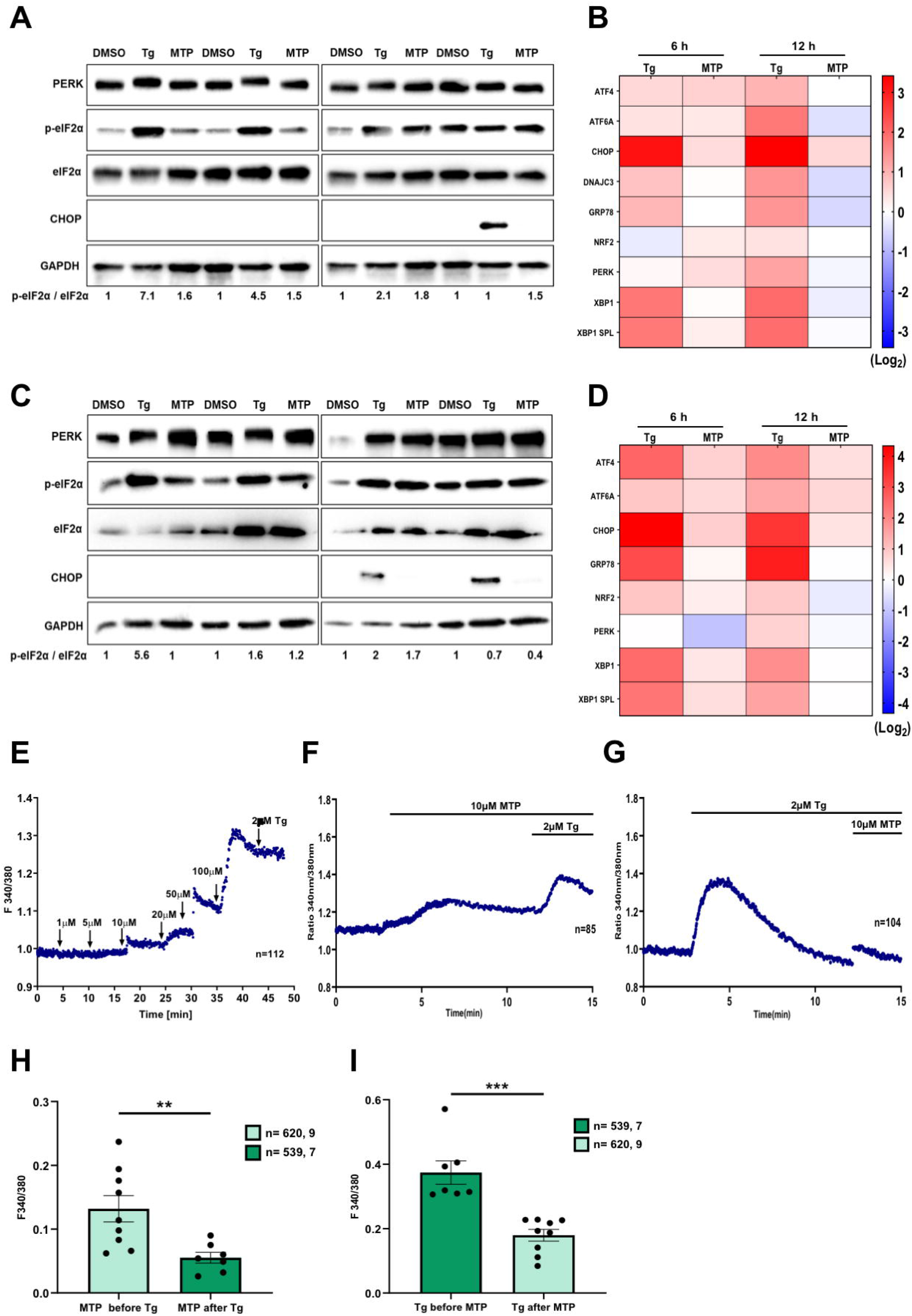
MTP activates adaptive ER stress and dysregulates ER calcium homeostasis. Neuro2a (A-B) and MEFs (C-D) were treated with DMSO/Tg (1 µM)/MTP (10 µM) for the indicated time points. (A & C) Protein lysates were immunoblotted for PERK, p-EIF2α, EIF2α, CHOP and GAPDH (loading control). The values below the blot show relative protein levels after normalization to DMSO controls. (B & D) mRNA levels of ER stress markers and chaperones were quantified using qRT-PCR. Heatmap depicts relative gene expression normalized to DMSO control, represented as mean (n=3). (E) Representative Ca^2+^ imaging trace of MTP dose response assay, where “n=112” denotes the number of cells in that particular trace. Cells were stimulated with increasing doses of MTP-1 μM, 5 μM, 10 μM, 20 μM, 50 μM and 100 μM followed by addition of 2 μM thapsigargin (Tg) in Ca^2+^-free buffer. (F) Representative Ca^2+^ imaging trace of experiments where cells were stimulated with 10 µM MTP in absence of extracellular Ca^2+^ followed by addition of 2 μM thapsigargin (Tg). Here, “n=85” denotes the number of cells in that particular trace. (G) Representative Ca^2+^ imaging trace of experiments where cells were stimulated first with 2 μM thapsigargin (Tg) to deplete ER Ca^2+^ stores, followed by addition of 10 µM MTP in absence of extracellular Ca^2+^. Here, “n=104” denotes the number of cells in that particular trace. (H) Quantitation of MTP (10 μM) induced ER Ca^2+^ stores depletion before and after the addition of 2 μM thapsigargin (Tg). 620 and 539 cells from 9 and 7 independent imaging dishes were analysed for the two conditions, respectively. (I) Quantitation of thapsigargin (Tg, 2 μM) induced ER Ca^2+^ stores depletion before and after the addition of 10 μM MTP. 539 and 620 cells from 7 and 9 independent imaging dishes were analysed for the two conditions, respectively. (“n11=11x, y” where “x” denotes total number of cells imaged and “y” denotes number of traces recorded). Data presented are mean ± S.E.M. Unpaired student’s t test, **, P<0.01; *** P < 0.001

MTP treated Neuro2a cells showed a modest enhancement of eIF2α phosphorylation which was consistent till 12 h (Fig 6A). These cells also showed activation of ATF4, CHOP, XBP1(s) and ATF6, but no activation of DNAJC, GRP78 & XBP1 at 6 h of treatment and downregulation of these transcripts by 12 h (Fig 6B). No CHOP protein expression was observed on MTP treatment (Fig 6A). The modest transcriptional activation of CHOP and absence of its protein expression on MTP treatment suggested the activation of an adaptive vs pro-apoptotic pathway of UPR induction.

Thapsigargin treated MEFs showed similar enhanced phosphorylation of eIF2α which attained baseline by 12 h (Fig 6C), and high transcriptional activation of the other ER stress reporter genes (Fig 6D). On the other hand, MTP treated MEFs showed marginal eIF2α phosphorylation, no detectable CHOP protein, and comparatively low transcriptional activation of the other genes (Fig 6C, D).

One of the most critical contributors to ER stress induction is dysregulated ER Ca^2+^ homeostasis. It is important to note that perturbations in ER Ca^2+^ signaling can lead to a variety of viral pathogenesis and therefore, it is emerging as a potential therapeutic target [30, 31]. Since MTP induces ER stress response, we examined if it can modulate ER Ca^2+^ signaling. We performed live cell Ca^2+^ imaging using ratiometric FURA-2AM dye in MEFs. The fluorescence intensity of FURA-2AM corresponds to cytosolic Ca^2+^ levels. We first performed a dose response assay with increasing concentration of MTP in absence of extracellular Ca^2+^. We observed that 10 µM MTP can induce an increase in cytosolic Ca^2+^ levels and 100 µM MTP completely depletes Thapsigargin (Tg) sensitive Ca^2+^ stores (Fig 6E). As these imaging assays were performed without Ca^2+^ in the extracellular bath, it suggests that the source of this rise in cytosolic Ca^2+^ levels is intracellular stores. ER is the major source of intracellular Ca^2+^ stores. Therefore, we examined if MTP is driving ER Ca^2+^ release to cytosol. We repeated the live cell Ca^2+^ imaging experiments with 10 µM MTP in absence of extracellular Ca^2+^ and observed a rise in cytosolic Ca^2+^ levels. In the same experiments, we then added Tg to block SERCA channels present on the ER, which led to further increase in cytosolic Ca^2+^ levels (Fig 6F). This suggests that MTP can only partially mobilize ER Ca^2+^ levels. We next performed opposite experiments wherein we first stimulated ER Ca^2+^ release with Tg and then gave MTP treatment (Fig 6G). If MTP induces Ca^2+^ movement from ER, then amplitude of this Ca^2+^ mobilization should be substantially decreased after Tg stimulation. Indeed, we observed that post Tg stimulation, MTP mediated cytosolic Ca^2+^ rise was significantly decreased (Fig 6G & H). Likewise, Tg induced ER Ca^2+^ release was drastically reduced upon pre-stimulation with MTP (Fig 6F & I). Taken together, these experiments demonstrate that MTP and Tg mobilize Ca^2+^ from same intracellular stores i.e. ER; most likely by acting over same target viz. SERCA channels. Further, these experiments establish MTP as a potent ER Ca^2+^ release inducer and that in turn at least partially explain the molecular mechanism connecting MTP treatment and induction of ER stress.

### MTP induces ER stress and negatively regulates type I interferon signaling in virus infected cells

Since MTP was a potent inducer of ER stress, we also examined the ER stress signatures in virus infected and drug treated cells. In agreement with our earlier study [17], JEV infected cells showed detectable eIF2α phosphorylation (Fig 7A), and upregulation of CHOP, GRP78, DNAJC3, ATF4, XBP1 & XBP1 spl. at 12 h of infection (Fig 7B). This corresponds to a stage where viral protein translation is well established in the cell. A combination of MTP treatment in JEV infected cells resulted in eIF2α activation at 6 h, a time point where virus replication complex biogenesis initiates, suggesting that MTP potentially exerts an inhibitory effect on early viral protein translation (Fig 7A). Also, ER stress transcripts were high in these cells at 3 hpi and persisted till later time points of infection. This suggests that the drug induces the activation of an adaptive ER stress response in these cells before infection is established. MTP treatment in LPS stimulated MEFs resulted in transcriptional activation of only ATF4, indicating an adaptive ER stress response (Fig S6A).

**Fig7:**
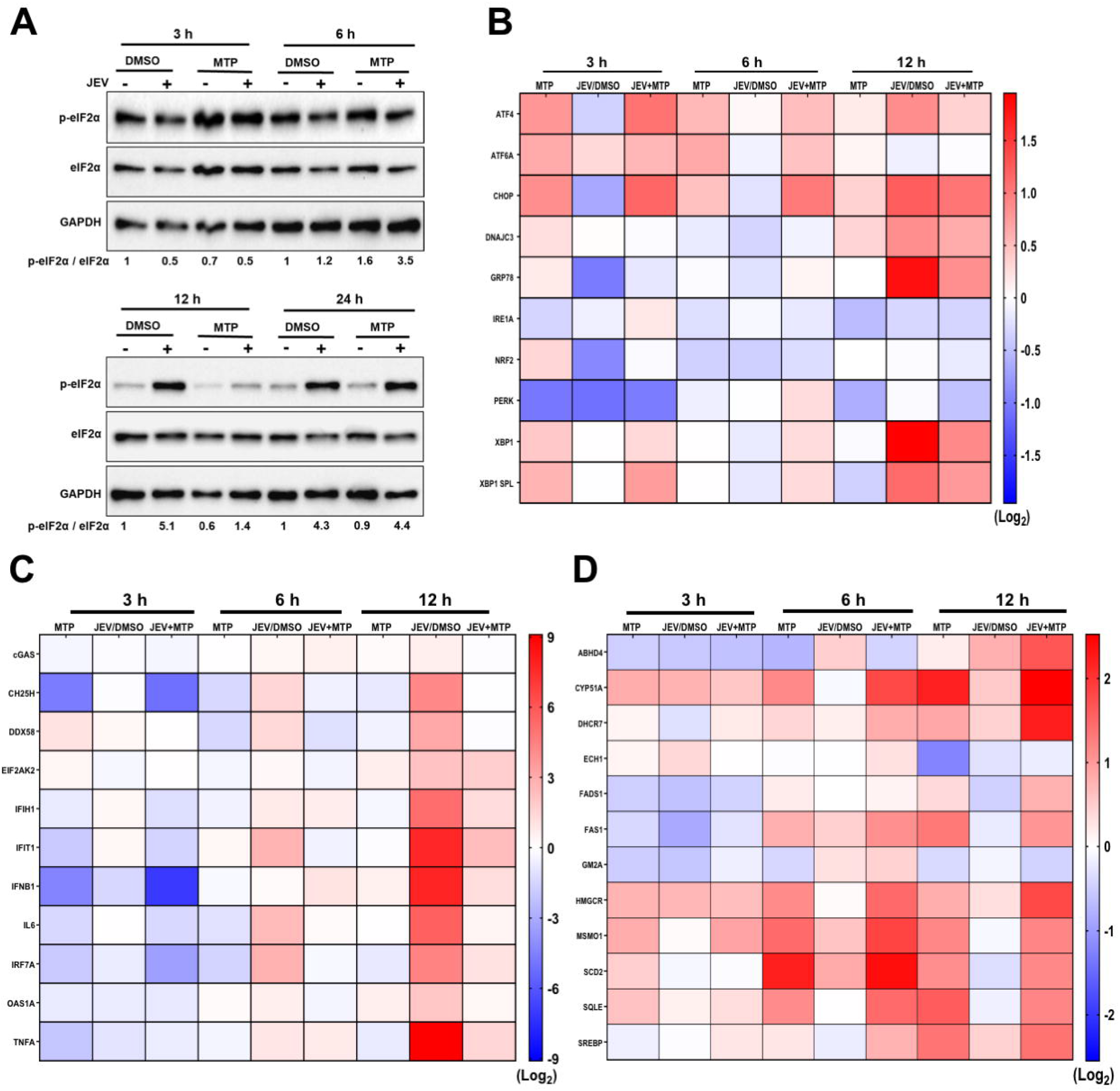
MTP induces adaptive ER stress and negatively regulates type I interferon signaling in virus infected cells. (A-D) MEFs were infected with JEV at MOI 1, 1 hpi, cells were treated with DMSO/MTP (10 µM) for the indicated time points. (A) Protein lysates were analyzed by immunoblotting using p-eIF2α, eIF2α, and GAPDH (loading control) antibodies. Values below blot show relative protein expression levels normalized to DMSO controls. (B-D) RNA was used for the quantitation of ER-stress (B), innate immune/inflammatory pathway (C), and cholesterol metabolic pathway genes (D). Heatmap showing relative gene expression levels normalized to DMSO control, represented as mean from two independent experiments.

Next, we examined the effect of MTP on type I interferon and inflammatory signaling in MEFs. MTP treatment of 3 h resulted in significant downregulation of basal transcript levels of IFIH1, IFIT1, IRF7, IFN-β, IL-6 and TNF-α (Fig7C). Interestingly, a highly significant reduction in transcript levels of cholesterol 25-hydroxylase (CH25H) were also observed starting at 3 h (Fig 7C). In JEV infected MEFs, innate immune and inflammatory transcripts showed upregulation starting 6 hpi, and were highly upregulated by 12 hpi (Fig 7C). In the virus and drug treatment group these were significantly reduced at all time points indicating that MTP exerts a strong negative impact on type I IFN and inflammatory responses in JEV infected cells (Fig 7C). A similar down-modulation was also observed in LPS+MTP treated MEFs suggesting that MTP treatment negatively regulates innate immune and inflammatory responses (Fig S6B).

CH25H is an IFN stimulated gene (ISG), that is upregulated in virus infection and is inversely related to cholesterol metabolic pathways in the cell. Further, the phenothiazines: clozapine and chlorpromazine have been shown to upregulate several genes of cholesterol and fatty biosynthesis, increase enzymatic activity of HMGCR, and enhance cholesterol and triglyceride levels in human glioma cells [32, 33]. We then checked the effect of MTP on genes of cholesterol metabolic pathways, and observed a significant upregulation of - Sqle, Cyp51A, SREBP, Hmgcr, Fas1, Msmo1, Scd2 (Fig 7D).

In JEV infected cells, no major effect on transcript levels of genes involved in cholesterol and lipid metabolic pathways was seen till 12 hpi, except for a slight downregulation of - Ech1 & Fads1 at 12 hpi. Interestingly, virus-infected drug-treated cells, showed higher transcript levels of SQLE, CYP51A, SREBP, Hmgcr, Msmo1, Fas1, Dhcr7, etc. suggesting that lipid/cholesterol metabolic pathways are highly upregulated in these cells (Fig 7F). Genes of lipid metabolic pathways were also significantly upregulated in LPS + drug treated cells (Fig S6C). It is possible that the observed downregulation of type I IFN and ISGs in MTP treated condition is directly linked to the activation of cholesterol metabolic pathways.

### Antiviral effect of phenothiazines is autophagy-dependent

We next attempted to elucidate which step of the virus life-cycle was being targeted by phenothiazines. We performed a time course analysis of virus infection in control and MTP/TFP treated Neuro2a cells (Fig 8A-B), and MTP treated MEFs (Fig 8C) and HeLa cells (Fig 8D). JEV life-cycle begins with a low endosomal pH mediated uncoating of the virus envelope, followed by nucleocapsid release into the cytosol, capsid dissociation, and translation of the plus-strand viral RNA into a single polyprotein that subsequently gives rise to virus structural and nonstructural proteins. Depending on the MOI and cell type this process takes 3-6 h, during which time the viral RNA levels decrease compared to 1 hpi, likely due to degradation of a fraction of the endocytosed virus in the endosomal system (Fig 8A-D). Once the viral RNA is translated, the virus replication complex is established on ER-derived membranes, and a rapid increase in virus replication is seen [34, 35]. A comparison of virus replication kinetics between control and drug treated cells clearly showed that the anti-viral effect was exerted at the level of viral protein translation/replication complex formation which was severely compromised in drug treated Neuro2a (Fig 8A-B), MEFs (Fig 8C) and HeLa cells (Fig 8D). Drug treatment also resulted in reduced transcript levels of various inflammation/cell death markers (Fig S7A), consistent with reduced cell death in drug treated infected cells.

**Fig8:**
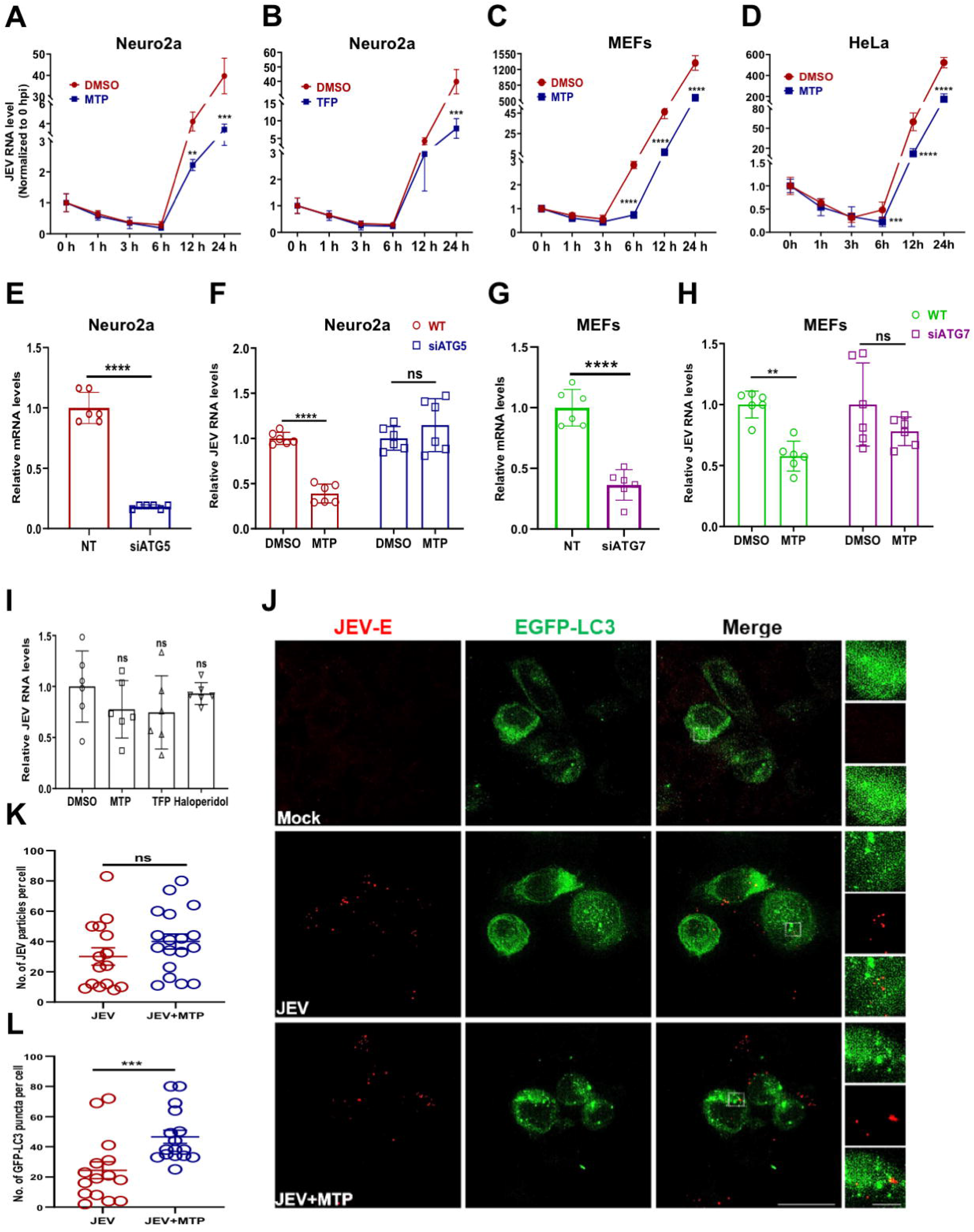
Antiviral effect of MTP is autophagy dependent. (A-D) Neuro2a (MOI 0.1), MEFs/HeLa (MOI 1) were mock/JEV infected and at 1 hpi, treated with DMSO/MTP (10 µM)/TFP (10 µM). Cells were harvested at the indicated hpi and viral RNA levels were quantified using qRT-PCR. Data represents values obtained from two independent experiments. (E) Neuro2a cells were transfected with siNT/siATG5 (50 nM) for 48 h, ATG5 RNA levels determined by qRT-PCR. (F) Neuro2a cells were transfected with siNT/ATG5 for 48 h and then infected with JEV (MOI 1). Cells were treated with 10 µM MTP at 1 hpi and harvested at 24 hpi to measure JEV RNA levels. Bar-graph shows relative viral RNA levels after normalization to respective NT transfected DMSO treated control. Data is plotted from two independent experiments (n=6). (G) MEFs were transfected with siNT/siATG7 (50 nM) for 48 h, bar-graph shows relative Atg7 RNA levels determined by qRT-PCR. (H) MEFs were transfected with siNT/ATG5 for 48 h, infected with JEV (MOI 1), and treated with 10 µM MTP at 1 hpi. JEV RNA levels were estimated at 24 hpi through qRT-PCR. Bar-graph shows relative viral RNA levels after normalization to siNT/DMSO control, data is plotted from two independent experiments (n=6). Data are represented as means ± SD. (I) Neuro2a cells were pre-treated with DMSO/MTP (10 µM)/TFP (10 µM)/Haloperidol (10 µM) for 1 h, and then infected with JEV (MOI 5) for 1 h in the presence of the drug. Cells were given trypsin treatment to remove dish/cell bound virus particles, and the levels of internalized virus were measured using qRT-PCR. Data is plotted from two independent experiments and compared by one-way ANOVA test. (J-L) Neuro2a cells stably expressing EGFP-LC3 were grown on glass coverslips and were mock/JEV (MOI 50) infected. At 1 hpi, cells were given DMSO/MTP treatment for 1 h. Cells were immunostained with JEV envelope antibody, and SIM imaging was performed. The right panel shows magnified view of the region marked by rectangle. Scale bar, 10 µm, 5 µm (inset). Graph shows quantitation of JEV particles (red dots) per cell (K), and autophagosome number (green puncta) per cell (L), calculated from 15-20 cells across two independent coverslips. Quantification was performed using Imaris 8 software and expressed as ± SEM. Statistical analysis was performed using unpaired student t-test, **, P<0.01; ***, P<0.001; **** P < 0.0001, ns; non-significant.

We next examined if the observed anti-viral effect of these drugs was autophagy dependent. siRNA-mediated depletion of Atg5 in Neuro2a cells (Fig 8E-F), and Atg7 in MEFs (Fig 8G-H) resulted in a complete loss of MTP anti-viral activity. Similar results were seen in ATG5 KO MEFs (Fig S7B-E) and ATG5 KO HeLa cells (Fig S7F-G). These data clearly indicated that the antiviral effect of the phenothiazines in diverse cell lines was mediated through autophagy.

JEV targets the dopaminergic neuron rich regions in the brain, and a D2R agonist has been shown to enhance infection, while the antagonists such as prochlorperazine, haloperidol and risperidone have been shown to suppress infection in neuronal cells [36, 37]. Both MTP and TFP did not lead to any reduction in virus entry (1 hpi) as measured through JEV RNA levels (Fig 8I), and through a quantitative immunofluorescence assay of JEV envelope antibody labelled virus particles in control and MTP treated cells (Fig 8J-K). As expected, the number of GFP-LC3 puncta were significantly increased in drug treated cells, however, no overlap of labelled virus particles with any autophagosome was observed (Fig 8J-L). These data suggested that the drug treatment does not inhibit virus entry, and the endocytosed virus particles do not appear to be targeted for virophagy.

### Anti-inflammatory effect of MTP in microglial cells is partially autophagy dependent

Microglial cells are crucial to modulate neuroinflammation through secretion of proinflammatory cytokines. We next checked if this process was autophagy dependent in the context of JEV infection (Fig S8). Secretion of proinflammatory cytokines IL-6, TNF-α, RANTES and MCP-1 from JEV infected microglial cells increased with increasing MOI (Fig S8B). Depletion of Atg5 in these cells resulted in significantly higher levels of cytokine secretion, indicating a crucial role of autophagy in mediating neuroinflammation (Fig S8A-B, 9A-B). Autophagy deficient microglial cells also displayed significantly enhanced JEV induced cell death (Fig S8C), however the JEV titers between autophagy competent and deficient microglial cells was not significantly different (Fig S8D).

We then checked if the anti-inflammatory effect exerted by MTP in microglial cells was autophagy dependent (Fig 9A). As shown earlier (Fig 1C-F, Fig 3D), MTP treatment resulted in significantly reduced levels of cytokine release from JEV infected cells (Fig 9B). MTP treatment of Atg5 depleted N9 cells resulted in reduced cytokine release however these levels were still significantly higher compared to MTP treated wild-type cells. These observations suggest that the anti-inflammatory effect of MTP on microglial cells cannot be attributed entirely to autophagy upregulation, and maybe mediated in part by MTP induced ER stress. A similar enhancement of ER stress (Fig S9A), and activation of lipid metabolic pathways (Fig S9B) was also seen in microglial cells. Collectively, these data indicate that MTP treatment displays an adaptive ER stress signature in microglial cells and exerts an anti-inflammatory effect that is partly autophagy dependent.

**Fig9:**
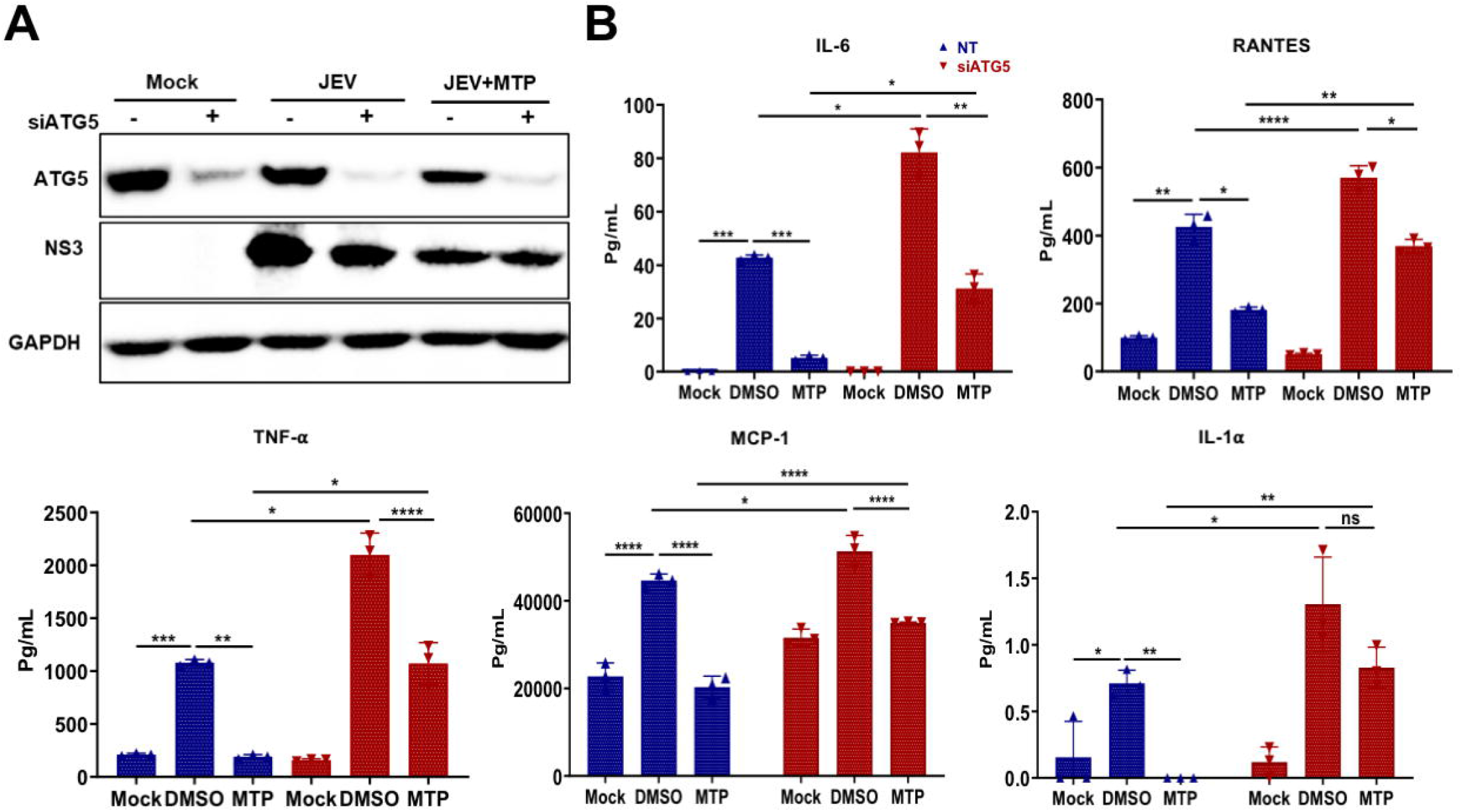
MTP inhibits inflammatory cytokines secretion from microglial cells partly through autophagy. N9 cells were transfected with siNT/ATG5 for 48 h, followed by mock/JEV (MOI 3) infection for 1 h and treatment with MTP (10 µM) till 24 hpi. (A) Cell lysates were prepared and proteins were analyzed by immunoblotting using ATG5, NS3 (infection control), and GAPDH (internal control) antibodies. (B) Culture supernatant was used for the quantitation of cytokine levels using flow cytometry-based CBA assay (n=3). Similar trends were seen in two independent experiments. All data were expressed as means ± SD and one-way ANOVA test was used to determine statistical significance. *, P<0.05; **, P<0.01; ***, P<0.001; **** P < 0.0001; ns; non-significant.

## Discussion

One-third of JE infections are fatal, and one-third develop permanent cognitive and/or motor defects due to severe neurological damage. The disease is acute and its pathogenesis is a combination of direct virus induced neuronal cell death and a massive neuroinflammatory response. Suppression of neuroinflammation in the patient is likely to be critical to improving prognosis, and this necessitates the need for development of effective therapies.

Our studies on JEV-host interactions have shown that infection-induced ER stress and autophagy are closely linked to virus replication and neuronal death [16, 17]. Several FDA approved drugs have been shown to enhance autophagy, and this provides a strong rationale for repurposing these drugs for treatment of diseases where autophagy upregulation could potentially provide a therapeutic benefit.

Based on published literature we curated a pool of FDA-approved drugs with autophagy-inducing potential and tested these for both antiviral and anti-inflammatory effects in vitro. Further, we investigated promising in vitro leads in the JE mouse model, at oral doses equivalent to the recommended human dose. The antipsychotic phenothiazine drug MTP showed significantly improved survival, reduced neuroinvasion and complete protection against BBB breach and neuroinflammation. MTP also known as Levomepromazine (brand name Nozinan) is prescribed for relief of moderate to severe pain and anxiety. Another widely prescribed antipsychotic phenothiazine, TFP showed similar antiviral and neuroprotective effects in vitro and in vivo. Our studies suggest that the antiviral and neuroprotective mechanism is likely to be a complex interplay of drug-induced dysregulation of Ca^2+^ homeostasis, adaptive ER stress, autophagy, downregulation of IFN/inflammatory response, and activation of cholesterol metabolic pathways. Indeed, it is challenging to establish a single/critical drug target, and establish a linear relationship in the sequence of events.

Drugs of the phenothiazine family are typical antipsychotics that are widely used in clinical practice for the treatment of bipolar disorders, psychosis and schizophrenia. The antipsychotic effect is attributed to the blockage of dopamine D2 receptors in the brain. The neuroprotective effects of these drugs in vitro and rodent models of Alzheimer’s, Parkinson’s, and Huntington’s disease are well documented [38–43], and has also been linked to autophagy induction [44]. Phenothiazine hydrochloride was first identified as an autophagy inducer in a high-throughput drug screening assay using a C. elegans model of protein aggregation [45]. Stress-dependent pharmacological activation of autophagy through TFP has been shown to have neuroprotective effects under conditions of α-synuclein accumulation in human dopaminergic neurons [46], and in Pink1 deficient zebrafish model and human cells [25].

Phenothiazines show diverse biological effects ranging from anti-cancer to anti-pathogen (virus, bacteria, fungus, protozoa) [47–50]. CPZ has shown antiviral effects against several viruses, including SARS-CoV-2 and flaviviruses such as JEV, DENV and WNV. Prochlorperazine has also shown antiviral activities against JEV, DENV, HCV and EBOV [36, 49, 51]. These drugs have been shown to alter cellular lipid dynamics [52], or obstruct endocytic pathways [53–56]. Besides acting as host-directed antivirals, phenothiazines have also been shown to directly interact with and destabilize the EBOV glycoprotein [57].

We observe that phenothiazines exert a strong antiviral effect at the level of viral protein translation/replication complex formation, which is dependent upon autophagy induction. While we observe no evidence of virophagy, the inhibition of virus replication could be either mediated by inhibition of viral protein translation or degradation of viral proteins in autolysosomes. MTP and TFP were potent autophagy inducers in several cell types and resulted in the activation of functional autophagy flux. Autophagy induction was observed to be mTOR independent and there was no TFEB translocation into the nucleus. MTP treatment also resulted in the transcriptional activation of genes involved in autophagosome expansion and formation (Atg12, Atg16l1, LC3A, LC3B).

The autophagy inducing properties of phenothiazines are documented though the mechanistic details lack clarity [22–24]. Studies in different cancer lines have shown both autophagy induction [24, 58-62], and autophagy flux inhibition [63–66]. CPZ is known to inhibit PI3K/Akt/mTOR in glioma and oral cancer cells [24, 62], and also induce ER stress in glioblastoma cell lines [60]. However, most studies showing phenothiazine induced mTOR inhibition and cancer cell cytotoxicity have used high drug doses in the range of 20-50 µM and beyond [24, 62, 67]. A structure-function relationship study identified a pharmacore that could induce neuronal autophagy in an Akt-and mTOR-independent manner. This was defined as the N^10^-substituted phenoxazine/phenothiazine, whereas the non-substituted phenoxazine and phenothiazine did not stimulate autophagy [68].

Our results demonstrate that phenothiazine induces adaptive ER stress and autophagy. One of the most critical signaling events that regulates both ER stress and autophagy is increase in intracellular Ca^2+^ levels [69, 70]. Therefore, we examined the role of MTP on cellular Ca^2+^ signaling and indeed, we found that MTP treatment results in rise in cytosolic Ca^2+^ levels. Our detailed live cell Ca^2+^ imaging experiments showed that the source of this increase in cytosolic Ca^2+^ is ER Ca^2+^ release. Since Tg (SERCA inhibitor) and MTP treatment showed a non-additive rise in Ca^2+^ levels, it suggests that they are mobilizing Ca^2+^ from same intracellular pools and most likely they act on same Ca^2+^ handling channel/pump. Indeed, literature suggests that a variety of phenothiazines including TFP can directly inhibit SERCA pump [71]. Taken together, our data suggests that similar to many other phenothiazines, MTP inhibits SERCA pump and induces an increase in cytosolic Ca^2+^ levels. Further, this rise in intracellular Ca^2+^ concentration can activate adaptive ER stress and autophagy. In future, it would be interesting to investigate precise molecular mechanism through which phenothiazines inhibit SERCA pumps as it would be relevant for several other disorders associated with SERCA hyperactivity.

MTP treatment resulted in rapid eIF2α phosphorylation and transcriptional activation of ATF4, CHOP, ATF6, Xbp1 and Xbp(s) indicating activation of ER stress. However, no CHOP protein was detectable in drug treated cells suggesting that the stress was primarily adaptive and not apoptotic. In cells exposed to ER stress, autophagy is transcriptionally activated as a survival response [27, 29, 72, 73]. Activation of CHOP is mediated by ATF4 and ATF6 [74, 75] and generally leads the cells towards apoptosis through upregulation of BH3 only proteins and downregulation of Bcl2 [28, 29]. CHOP also drives inflammation through secretion of IL-1β through caspase-11/caspase-1 [76], and activation of NFkB [77], and can negatively impact cholesterol and lipid biosynthesis pathways [78]. Studies have shown that in response to aa starvation/tunicamycin treatment, the autophagy genes Atg16l1, Mapi1lc3b, Atg12, Atg3, Beclin1 and Gabarapl2, can be activated by ATF4 independently of CHOP [26, 79]. The autophagy gene activation profile induced by MTP in our study was also similar, along with upregulation of the pro-survival gene Bcl2. Our data indicates that MTP induces a unique chronic/adaptive ER stress with gene expression profiles that are qualitatively distinct from those induced by severe stress such as Thapsigargin. This unique adaptive ER stress signature that also includes transient phosphorylation of eIF2α, is likely to contribute to an antiviral state.

The phenothiazines have been shown to inhibit cytokine secretion from microglial cells in animal models of traumatic brain injury, subarachnoid/intracerebral hemorrhage, hypoxia-ischemic recovery etc. [80–87]. Patients with schizophrenia and first episode psychosis (FEP) display abnormal profiles of proinflammatory cytokines (especially IL-6 and TNF-α) prior to start of treatment. In these individuals, antipsychotic treatment resulted in decreased serum concentrations of IL-1β, IL-6, IFN-γ, TNF-α, and showed therapeutic effects by reducing microglial inflammation comparable to levels in healthy controls [88]. Indeed, we observed a similar effect of the drugs in inflammatory cytokine release from both microglial cells and BMDMs. This effect was partially reversed by autophagy inhibition, suggesting that mechanisms other than autophagy could also be contributing to the anti-inflammatory effect of the phenothiazines.

TFP and fluphenazine have been shown to be direct inhibitors of TLR3-IRF3 signaling pathway [89]. In our study MTP also caused significant inhibition of type I IFN and several other ISGs, indicating suppression of innate immune responses. This was not due to reduced virus replication, as a similar inhibition was also seen in response to LPS treatment. While type I IFN is primarily antiviral in nature, it might also exert a proinflammatory function. Interestingly, we also observed significant downregulation of CH25H gene, which links lipid and cholesterol homeostasis to inflammation and immune response [90, 91]. While the CH25H gene product, 25-hydroxycholesterol, has primarily an antiviral function [92], it also acts as an amplifier of inflammatory signaling in macrophages and enhances tissue damage [91].

It is also well established that several antipsychotics can cause metabolic disorders such as hypertriglyceridemia, glucose dysregulation and elevated cholesterol levels, which is attributed to transcriptional activation of cholesterol and fatty acid biosynthetic genes via SREBP1 and SREBP2 [32, 33, 93, 94]. We too observed that MTP treatment resulted in transcriptional upregulation of several cholesterol biosynthetic genes in divergent cell types. An inverse relation between type I IFN response and flux through the mevalonate pathway has been reported earlier [95, 96]. MTP mediated enhancement of sterol biosynthesis could be directly responsible for the observed downmodulation of type I IFN and inflammation, however this requires further validation.

In conclusion, our study provides evidence that the phenothiazines MTP and TFP have robust antiviral and neuroprotective effects in JE disease condition and have the potential to be repurposed for treatment. These drugs are approved for chronic use and have a high therapeutic index. As a future scope of this work, a small investigator-initiated clinical trial of this drug in JEV patients can test the findings of this pre-clinical study and establish proof of concept in humans.

## Materials and Methods

### Ethics Statement

All animal experiments were approved by the Institutional Animal Ethics Committee of the Regional Centre for Biotechnology (RCB/IAEC/2018/039). Experiments were performed as per the guidelines of the Committee for the Purpose of Control and Supervision of Experiments on Animals (CPCSEA), Government of India.

### Cell lines and virus

Neuro2a (mouse neuroblastoma), C6/36 (insect), and Vero cell lines were obtained from the cell repository at the National Centre for Cell Sciences Pune, India. WT and atg5−/− Mouse embryonic fibroblasts (MEFs) were obtained through the RIKEN Bio-Resource Cell Bank (RCB2710 and RCB2711). WT and atg5−/− HeLa cell lines were a kind gift from Dr. Richard J. Youle (NIH, USA); and mouse microglia N9 cell line was a gift from Prof. Anirban Basu (NBRC, India).

Neuro2a cells stably expressing EGFP-TFEB/EGFP-LC3 were generated through plasmid transfection, growth in G418 selection media followed by single cell isolation through FACS. MEFs stably expressing GFP-LC3-RFP-LC3ΔG [97], were generated through retroviral transduction as described earlier [98].

Dulbecco’s modified Eagle’s medium (DMEM) was used to culture Neuro2a, MEFs and HeLa cells, Eagle’s minimal essential medium (MEM) for Vero cells, RPMI for N9 cells, and Leibovitz’s (L-15) medium was used to culture C6/36 cells. All media were additionally supplemented with 10% fetal bovine serum (FBS), 100 μg/ml penicillin-streptomycin, and 2 mM L-glutamine.

JEV isolate Vellore P20778 strain (GenBank accession no. AF080251) was generated in C6/36 cell line. UV-inactivated JEV was generated by exposure of virus to UV (1600 x 100 µ J/cm^2^) for 20 min on ice. For animal experiments the mouse-adapted JEV-S3 strain was used [21]. Virus titration was performed in Vero cells using plaque assays as described earlier [35]. All reagents, antibodies and plasmids used in the study are listed in Table S2.

### Primary cell culture

#### Bone marrow-derived macrophage (BMDMs)

BMDMs were isolated from 6-7 weeks old C57BL/6 mice. Briefly, mice were euthanized and femurs were dissected, washed with PBS and RPMI media, and flushed with L929-conditioned medium to extrude bone marrow. After RBC lysis, cells were cultured in RPMI complete media supplemented with L929-conditioned media for 7 days. BMDMs were detached using 10 mM EDTA and seeded in 24 well plates for virus infection and drug treatment experiments.

#### Embryonic cortical neurons

Mouse primary cortical neuronal cells were isolated from pregnant mice at embryonic day 16.5 (E16.5). Briefly, embryos were collected by decapitation from pregnant mice, the cortices were dissected from isolated embryonic brains and collected in dissociation media, HBSS (1X sodium pyruvate, 20% glucose, 1 M HEPES, pH 7.3). Tissues were digested with trypsin and DNAse I to make single-cell suspensions. Cells were washed, centrifuged, and resuspended in complete neurobasal medium supplemented with 10% FBS, 20% glucose, 1X Sodium pyruvate, and antibiotics. Finally, cortical neuronal cells were plated on poly-l-lysine coated plates, and media was changed with maintenance media (neurobasal B-27, 1X glutamine, penicillin-streptomycin solution) after every 2 days by adding half new maintenance media.

### Virus infection and cell treatment

All virus infection studies were performed by giving mock/JEV infection at indicated MOI for 1 h. Cells were then washed and complete medium was added. Drug treatment was given by adding 10 µM drug/DMSO (control) to mock/JEV-infected cells, which was maintained till the end of the experiment as indicated. Cells were harvested for cell viability assays, RNA isolation or western blotting. Culture supernatants were used for quantitative estimation of cytokines using Cytokines bead array (CBA), or virus titers through plaque assay. For autophagy flux and lysosome pH assays, cells were treated with DMSO/Torin1 (1 µM)/MTP (10 µM) /TFP (10 µM) for 6 h. siRNA treatment was performed using mouse-specific Atg5/Atg7/non-targeting (NT) siRNA (50 nM, ON-TARGET plus SMART pool) using the transfection reagent Dharmafect 2 for Neuro2a cells, and Lipofectamine^TM^ RNAimax for MEFs and N9 cells. At 48 h post-transfection, cells were harvested and the knockdown efficiency was measured using western blotting or qRT-PCR. LPS treatment was given at 1 µg/ml for indicated times. Every experiment had biological triplicates and was performed two or more times.

### RNA isolation and Quantitative Real Time (qRT)-PCR

RNA was extracted using Trizol reagent. cDNA was prepared using ImProm-II™ Reverse Transcription System kit, and used to set up qRT-PCR on the QuantStudio 6 (Applied Biosystems). JEV RNA level was determined by specific Taqman probes, and GAPDH was used as an internal control. The gene expression for autophagy, UPR, innate immunity, and lipid/cholesterol biosynthesis was performed with SYBR green reagents. The expression of each gene was calculated by normalization to respective mock/DMSO controls. Each experiment had biological triplicates, and qPCR for each sample was done in technical triplicates. The primer sequences for all the genes tested in the study are listed in Table S3.

### Western blotting

Cells were lysed in cell lysis buffer (150 mM NaCl, 1% Triton X-100, 50 mM Tris-HCl pH 7.5, 1 mM PMSF, and protease inhibitor cocktail) for 45 min at 4°C. The supernatant was used for the estimation of protein concentration using BCA assay kit. Cell lysates were mixed with 4X Laemmli buffer (40% glycerol, 20% β-mercaptoethanol, 0.04% bromophenol blue, 6% SDS, 0.25 M Tris-HCl pH 6.8) and boiled at 95°C for 10 min to denature proteins. Equal concentrations of cell lysates were separated by SDS-PAGE and transferred to PVDF membranes for immunoblotting. The blots were visualized using a Gel Doc XR+ gel documentation system (Bio-Rad) and the expression of proteins was calculated by measuring the intensity of bands using ImageJ (NIH, USA) software. The fold change was calculated after normalization with respective loading controls. All western blotting experiments were performed three or more times, and representative blots are shown.

### Cytokines bead array (CBA)

Mouse microglia N9 cells (biological triplicates) were mock/JEV-infected at indicated MOIs for 1 h. At 12 hpi, cells were treated with DMSO/drugs for 24 h. Alternately N9 cells were treated with 1 µg/ml LPS for 24 h. Supernatants were harvested and used to quantitate the levels of cytokine IL-6, TNF-α, MCP-1, RANTES, IL-1β, and IL-1α using CBA assay as per manufacturer’s instruction. The analysis was performed with FCPA array software and the concentration of each cytokine was determined based on their standard curve. All CBA assays were performed two or more times, and representative data from one experiment is shown.

### Animal experiments

The mouse adapted strain JEV-S3 was generated in 3-4 day old C57BL/6 mice pups as described earlier [21]. Briefly, pups were infected with JEV (10^5^ PFU) by an intracranial route. By 3-4 dpi, symptoms of JEV infection such as movement impairment and constant shivering/body tremors were observed. The pups were sacrificed, and the brain tissues were harvested, homogenized in incomplete MEM media, and the supernatant containing infectious virus was titrated by plaque assay.

For JEV survival and other experiments, 3-week-old C57BL/6 mice of either sex were weighed and randomly divided into mock/drug/JEV/JEV+drug groups. Mice were infected intraperitoneally with 10^7^ PFU JEV, while the mock-infected group received equal volume of incomplete DMEM media by intraperitoneal injection. At 4 hpi, mice were treated with vehicle control (PEG 400) or the drugs MTP (2 mg/kg)/Flubendazole (5 mg/kg)/Fluoxetine (5 mg/kg)/Rilmenidine (5 mg/kg)/TFP (1 mg/kg) in PEG 400 formulations by oral route every 24 h till 15 days. Mice were monitored for symptoms of JEV infection such as change in body weight, movement restrictions, piloerection, tremor, body stiffening, hind limb paralysis, and mortality.

To determine cytokine levels in brain, mice (n=3) were sacrificed from each group on days 3, 4, 5, 6, and 7, and brain tissues were collected. These were homogenized in lysis buffer, and 30 μg of protein from each sample was used for quantitation of cytokines using LEGENDPLEX MU anti-virus response panel 13-plex assay kit as per manufacturer’s instructions. Data was analyzed with LEGENDplex^TM^ Multiplex assay software and the concentration of each cytokine was calculated based on their standard curve.

#### Evans blue leakage assay

Mice were intra-peritoneally injected with 100 µl of 2% Evans blue at 3 and 6 dpi. After 45 min they were sacrificed and brains were harvested. Images were captured to visualize the distribution of dye in the brain. For dye quantification, the tissues were weighed and homogenized in dimethylfumarate (DMF) (200 mg/500 μl DMF). The homogenate was heated at 60°C overnight to ensure complete extraction of the dye. The samples were then centrifuged and absorbance of each sample was measured at 620 nm. Evans blue content was quantitated using a standard curve.

### Cell viability assays

Cell viability assays were performed using the CellTiter-Glo® assay kit as per manufacturer’s instructions. The percentage of cell viability was calculated as: [(ATP luminescence for experimental condition) / (ATP luminescence for untreated condition)] X 100 and normalized to mock-infected/untreated DMSO treated control. MTT assay was performed as described earlier [35]. LDH assay using culture supernatant from siNT/Atg5 treated microglial cells was performed using CyQUANT™ LDH Cytotoxicity Assay kit as per manufacturer’s instructions. The percentage cell death was normalized to siNT mock-infected control.

### Immunofluorescence studies

#### Image-based high content screening for virus replication complex

Neuro2a cells were seeded in 96 well-black polystyrene microplates (Corning, CLS3603), and infected with JEV at MOI 5. At 1 hpi cells were treated with DMSO/drugs for 24 h, fixed with 4% paraformaldehyde (PFA) and permeabilized with 0.3% Tween-20 for 30 min at RT. After blocking with 5% BSA, JEV-NS1 primary antibody was added (1 h), followed by Alexa Fluor-labelled specific secondary antibody (1 h), and final incubation with DAPI (0.5 µg/ml) for 15 min. Images were acquired from the entire well area (16 fields per well) on ImageXpress Micro Confocal High-Content Imaging System (Molecular Devices, USA) using FITC and DAPI channels with a 10 X objective lens. The percentage of JEV-NS1 positive cells was calculated using the multi-wavelength cell scoring module of the MetaXpress software.

#### TFEB nuclear translocation assay

EGFP-TFEB stable Neuo2a cells were treated with DMSO/Torin1 (1 µM)/MTP (10 µM) /TFP (10 µM) for 6 h, fixed, and stained with DAPI, and images were acquired as described above. The Pearson coefficient between GFP and DAPI was calculated using the translocation module of the MetaXpress software (cut off =0.5). The percentage of EGFP-TFEB nuclear translocation per well was calculated using Torin1 as a positive control.

#### Autophagy flux assay

The flux assay was performed as described earlier [98]. MEFs stably expressing GFP-LC3-RFP-LC3ΔG were seeded at a density of 10,000 cells/well in 96-well black polystyrene microplates, followed by treatment of DMSO/Torin1 (1 µM)/Baf A1 (100 nM)/MTP (10 µM)/TFP (10 µM) for 6 h. Cells were fixed with 4% PFA and stained with DAPI. Images were recorded from 16 fields per well that covered the entire well area, on ImageXpress Micro Confocal High-Content Imaging System using DAPI, FITC, and Texas red channels with a 10 X objective lens. Analysis of images was performed using multi-wavelength cell scoring module of the MetaXpress software, that calculates the integral intensity of GFP and RFP from triple positive (DAPI, GFP, RFP) cells. GFP/RFP ratio was estimated and normalized to DMSO-treated control. For autophagy flux inducer, the cut-off value of GFP/RFP was set to < 0.8, while GFP/RFP > 1.2 was considered for autophagy flux inhibitors. Torin1 (GFP/RFP= 0.59) and BafA1 (GFP/RFP= 1.25) were used as positive and negative controls respectively.

#### GFP-LC3 puncta formation

Neuro2a cells stably expressing EGFP-LC3 were grown on glass coverslips. These were treated with DMSO/Torin1 (1 µM)/ MTP (10 µM) /TFP (10 µM) for 6 h, fixed with 4% PFA and mounted using ProLong Gold anti-fade reagent with DAPI. Images were acquired by Elyra PS1 (Carl Zeiss Super-resolution microscope) with 60X objective (lasers 405, 488 nm). LC3 puncta were counted from 20 cells acquired from two independent coverslips using ‘Analyse particles’ plugin algorithm of ImageJ (Fiji).

### Endosome Acidification Assay

LysoTracker Red and LysoSensor Yellow-Blue assays were performed as described earlier [98, 99] Briefly, Neuro2a cells were grown on glass coverslips, and treated with DMSO (control)/Torin1 (1 µM)/ BafA1 (100 nM) or MTP/TFP (10 µM) for 6 h, followed by incubation with 10 µM LysoTracker red (40 min), or 10 µM LysoSensor Yellow-Blue (5 min). Cells were then washed with ice-cold PBS three times and fixed using 4% PFA. Imaging was done on LSM 880 microscope, Carl Zeiss. LysoSensor Yellow-Blue imaging was done using the excitation wavelength range of 371-405 nm and emission wavelength range of 420-650 nm. The LysoSensor dye has dual-emission peaks of 440 nm (blue in less acidic organelles) and 540 nm (yellow in more acidic organelles). The analysis of LysoTracker red, and LysoSensor Yellow-Blue (yellow) fluorescence intensities were performed using ImageJ (Fiji) and normalized to DMSO-treated control, from 50 cells across two independent coverslips.

### Measurement of oxidative stress

Neuro2a cells were mock/JEV (MOI 5) infected for 1 h, followed by DMSO/FDA drug (10 µM) treatment till 24 hpi. DMF (70 µM) and NAC (3 mM) were added at 16 hpi and maintained till 24 hpi. Post-treatment, cells were incubated with 5 mM of oxidative stress indicator CM-H2DCFDA in incomplete media for 15 min. Cells were washed with PBS and fluorescence intensity was measured using flow cytometry BD FACS Verse (BD Biosciences, USA). All FCS files were analysed via FlowJo software and represented as mean fluorescence intensity.

### Virus entry assay through qRT-PCR

JEV entry can be quantitatively measured by estimating endocytosed virus levels through qRT-PCR at 1 hpi as described previously [35]. Briefly, Neuro2a cells were pre-treated with DMSO/MTP (10 µM) /TFP (10 µM) for 1 h, and then infected with JEV (MOI 5) for 1 h in the presence of the drug. Cells were harvested by washing, and trypsin treatment was given to remove any extracellular attached virus. qRT-PCR was done to measure the levels of internalized viral RNA relative to the GAPDH transcript.

### Immunofluoresence based virus entry assay

Neuro2a cells stably expressing EGFP-LC3 were grown on glass coverslips. These were mock/JEV (MOI 50) infected, and DMSO/MTP (10 µM) was added at 1 hpi and maintained for another 1 h. The cells were then fixed with 4% PFA and immunostained for the JEV envelope antibody. Coverslips were mounted using ProLong Gold anti-fade reagent with DAPI, and images were acquired by Elyra PS1 (Carl Zeiss Super-resolution microscope) with 63 X objective (lasers 405, 488 and 561nm). Colocalization between EGFP-LC3 and JEV-envelope was calculated by individual Spots and Spot to Spot colocalization per cell using Imaris 8 software.

### Calcium Imaging

Calcium imaging was performed as reported earlier [100, 101]. MEFs were cultured on confocal dishes (SPL life sciences, 200350) to attain 60-80% confluency. Cells were incubated in culture medium containing 4 μM fura-2AM for 45 min at 37°C. Post-incubations, cells were washed 3 times and bathed in HEPES-buffered saline solution (2 mM CaCl2, 1.13 mM MgCl2, 140 mM NaCl, 10 mM D-glucose, 4.7 mM KCl and 10 mM HEPES; pH 7.4) for 5 min. Further, 3 washes were given and cells were bathed in HEPES-buffered saline solution without 2 mM CaCl2 to ensure removal of extracellular Ca^2+^ before starting the measurements. A digital fluorescence imaging system (Nikon Eclipse Ti2 microscope coupled with CoolLED pE-340 Fura light source and a high-speed PCO camera) was used, and fluorescence images of several cells were recorded and analyzed. Excitation wavelengths-340nm and 380nm were used alternately for Fura-2AM and emission signal was recorded at 510nm. Ca^2+^ traces represent average data from multiple cells from a single imaging dish (number of cells is denoted by “n” on the graphs). Bar graphs represent data from multiple imaging experiments. The number of cells and traces for different conditions are specified on the respective bars as n=x, y where x stands total number of cells imaged and y means total number of independent experiments performed.

### Statistical analysis

Statistical analysis was performed using paired/unpaired Student’s t-test, one/two-way ANOVA test, and Log-rank (Matel-Cox) test. Differences were considered significant at P values of *p <0.05, **p <0.01, ***p <0.001, and ****p <0.0001, as indicated in the figures. Error bar indicates means ± SD/SEM. All graphs were plotted and analysed using GraphPad Prism 8.

### Data Availability Statement

This study includes no data deposited in external repositories.

## Acknowledgements

We thank Padmakar Tambare and the staff of the Small Animal Facility for their help with animal experiments. We also acknowledge facilities and staff of Advanced Technology Platform Centre. We are grateful to Dr. Kiran Bala Sharma for critical inputs on the manuscript. All Virology lab members are acknowledged for useful discussions and support.

## Author contributions

Experiments & data analysis (SKP, LM, SK, SO, KA, PS, RS); methodology (SKP, LM, SK, KA, PS, AB, RKM, SV, MK); formal analysis (SKP, LM, KA, DM, AB, RKM, SV, MK); resources (AB, RM, SV, MK); writing (SKP, LM, KA, RKM, MK); funding & project administration (MK).

