## Supplementary Data for "Methotrimeprazine exerts antiviral and neuroprotective effects in Japanese encephalitis virus infection through activation of adaptive ER stress and autophagy"

#### Supplementary Figure Legends

**FigS1: Effect of FDA-approved drugs on JEV-induced neuronal cell death.** (A) Neuro2a cells were treated with a panel of 42 FDA-approved drugs at a concentration of 10 µM for 48 h, and cell viability was measured through ATP luminescence using CellTiter-Glo® assay kit. The percentage cell viability was calculated after normalization to DMSO-treated control (n=3). (B) Neuro2a cells were mock/JEV/UV-inactivated JEV infected at MOI 1, 5 and 10. Cells were harvested at 24, 48, and 72 h. (C) Neuro2a cells were infected with JEV at MOI 5, and at 2 hpi treated with indicated drugs (10 µM) till 48 hpi. (B-C) The percentage cell viability was calculated by normalization to mock-infected control. Data are expressed as means ± SD from biological triplicates. Statistical analysis was performed using one-way ANOVA. *, P<0.05; **, P<0.01; ***, P<0.001; ****, P<0.0001.

**FigS2: Effect of FDA-approved drugs on JEV-induced inflammatory response.** (A) N9 cells were infected with JEV (MOI 1) and harvested at indicated time-points. Relative viral RNA levels were measured by qRT-PCR (n=3). (B-E) N9 cells were mock/JEV/UV-JEV infected (MOI 1) and cytokine levels were quantitated at the indicated time points using CBA assay (n=3). (F) N9 cells were treated with indicated drugs (10 µM) for 24 h, the percentage cell viability normalized to DMSO control was calculated (n=3). (G) N9 cells were mock/JEV (MOI 1) infected, and at 1 hpi treated with DMSO/drugs (10 µM) till 24 hpi. Relative JEV RNA levels were determined using qRT-PCR. Data is represented from three independent experiments. (D) N9 cells were treated with DMSO/LPS (1 µg/ml)/LPS+MTP (10 µM) for 24 h, and cytokine levels were quantified from the supernatant (n=3). One-way ANOVA test was used to determine statistical significance. *, P<0.05; ***, P<0.001.

**FigS3: Antiviral effect of shortlisted drugs in JEV-mouse model.** C57BL/6 (3 weeks old) mice were mock/JEV-S3 ($10^7$ pfu) infected through an i.p. injection, and at 4 hpi, treated with vehicle control (PEG400)/Flubendazole (5 mg/kg)/Fluoxetine (5 mg/kg)/MTP (2 mg/kg)/Rilmenidine (5 mg/kg) by oral gavage at an interval of 24 h till 15 days. Survival curve of mock (n=4)/ Flubendazole (n=4)/ Fluoxetine (n=4)/ MTP (n=4)/ Rilmenidine (n=4)/ JEV (n=7)/ JEV+flubendazole (n=7)/

JEV+fluoxetine (n=7)/ JEV+MTP (n=7)/ JEV+Rilmenidine (n=7) was plotted, a Log-rank (Mantel-Cox) test was used to determine the statistical significance comparing vehicle and drug-treated infected mice group. *, P<0.05.

**FigS4: MTP inhibits inflammatory cytokines secretion in response to proinflammatory stimuli.** BMDMs were treated with DMSO/MTP (10 µM)/LPS (1 µg/ml)/LPS+MTP for 24 h, supernatant was harvested and used for the quantitation of cytokine levels (n=3). All data were expressed as means ± SD and statistical significance was determined using one-way ANOVA test, **** P < 0.0001, ns; non-significant.

**FigS5: Phenothiazines induce functional autophagy flux and do not alter lysosomal pH.** (A-B) EGFP-LC3 expressing stable Neuro2a cells were treated with DMSO/Torin1 (1 µM)/MTP (10 µM) /TFP (10 µM) for 6 h. (A) Representative SIM images are shown. Scale bar, 10 µm. (B) Bar-graph shows quantitation of EGFP-LC3 puncta per cell. Data is acquired from 20 cells across two independent coverslips. (C-D) GFP-LC3-RFP-LC3ΔG expressing stable MEFs were treated with DMSO (control)/Torin1 (1 µM)/ BafA1 (100 nM) or MTP/TFP (10 µM) for 6 h. (C) Images were visualized by high-content imaging system. Scale bar, 100 µm. (D) Graph showing GFP/RFP ratios. (E-H) Neuro2a cells grown on glass coverslips were treated with indicated drugs as described above for 6 h, followed by incubation with 10 µM LysoTracker Red for 40 min (E-F) or 10 µM LysoSensor Yellow-Blue for 5 min (G-H). Representative confocal images are shown. Scale bar, 10 µm (E); 20 µm (G). LysoTracker Red (F) and LysoSensor Yellow-Blue (yellow) (H) fluorescence intensities were calculated from 50 cells across two independent coverslips using ImageJ (Fiji). All data are normalized to DMSO control and expressed as means ± SD, one-way ANOVA test, *, P<0.05; **, P<0.01; **** P < 0.0001.

**FigS6: MTP induces adaptive ER stress and negatively regulates type I interferon signaling in LPS treated cells.** (A-C) MEFs were treated with MTP (10 µM)/LPS (1 µg/ml)/LPS+MTP for 6 h, and relative mRNA transcript levels of ER stress (A), innate immunity (B) and cholesterol metabolic genes (C) were determined by qRT-PCR. Data are represented as means ± SD (n=3).

**FigS7: Antiviral effect of phenothiazines is autophagy dependent.** (A) Mock/JEV (1 MOI) infected MEFs were treated at 1 hpi with MTP (10 µM) for 12/24 h and mRNA levels of cell death genes was determined by qRT-PCR. The heatmap shows relative gene expression level after normalization to DMSO treated control. (B-G) WT and ATG5-/- MEFs (B, D) and WT and ATG5-/- HeLa cells (F), were treated with indicated concentrations of MTP/TFP for 24 h, and the percentage cell viability was measured and normalized to respective DMSO treated controls. (C, E & G) WT and ATG5-/- MEFs/HeLa cells were infected with JEV at MOI 1, and at 1 hpi treated with MTP/TFP at indicated concentrations. Cells were harvested at 24 hpi and the relative viral RNA levels were quantitated using qRT-PCR, and plotted after normalization to respective DMSO-treated control. Data represents values from three independent experiments, expressed as means ± SD. One-way ANOVA test was used for the determination of statistical significance, **, $P<0.01$; ***, $P<0.001$; **** $P < 0.0001$, ns; non-significant.

**FigS8: Autophagy restricts JEV-induced inflammatory response in microglial cells** (A) N9 cells were transfected with siNT/ATG5 for 48 h, followed by mock/JEV infection (1/ 3/ 5 MOI) for 24 h. (A) Western blotting was performed using ATG5, NS3 (infection control), and GAPDH (internal control) antibodies. (B) Quantification of cytokine levels from the culture supernatant. (C) JEV-induced cell death determined through LDH release assay. (D) JEV titres. (n=3), unpaired student t-test, *, $P<0.05$; **, $P<0.01$; ***, $P<0.001$; **** $P < 0.0001$, ns; non-significant.

**FigS9: MTP activates ER stress and cholesterol metabolic pathway genes in microglial cells.** N9 cells were mock/JEV (MOI 1) infected, and at 1 hpi treated with DMSO/MTP (10 µM) till 24 hpi. RNA levels of the indicated genes: ER stress panel (A), and cholesterol metabolic pathways (B) was determined by qRT-PCR. Heatmap shows relative gene transcript levels normalized to DMSO controls.

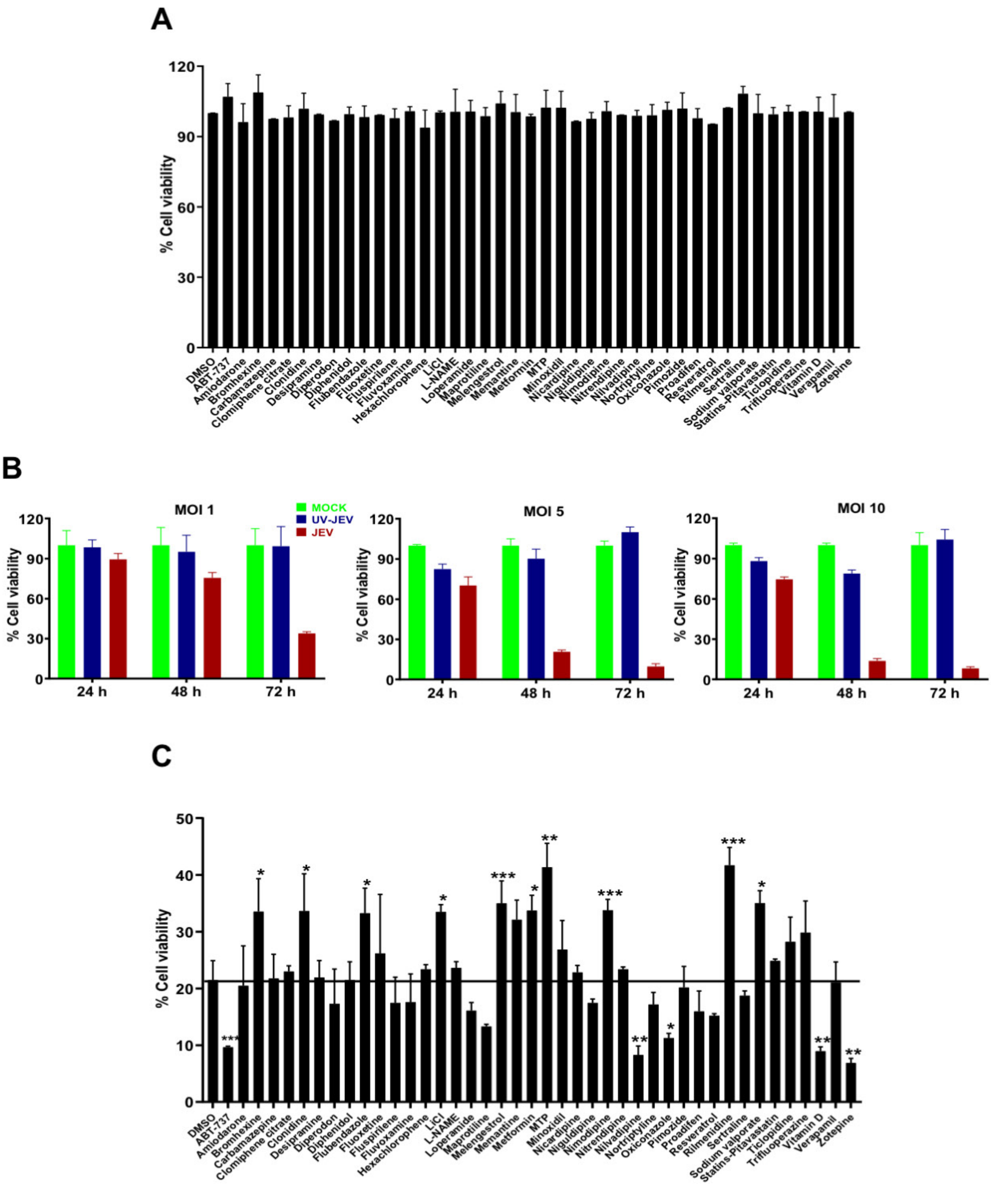
A
% Cell viability
B
MOI 1
MOI 5
MOI 10
MOCK
UV-JEV
JEV
24 h
48 h
72 h
C

Fig.S1

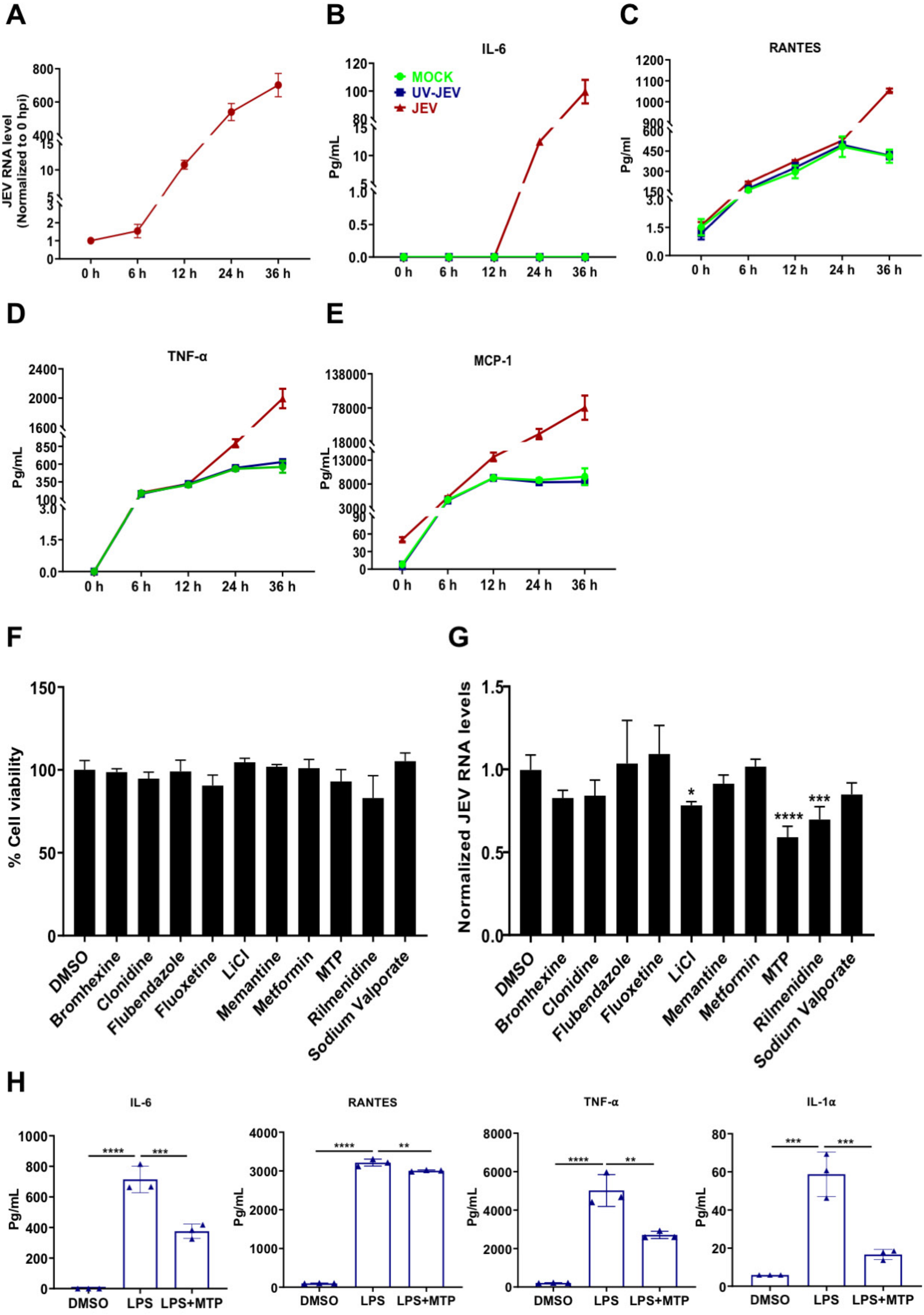

A
JEV RNA level (Normalized to 0 hpi)
B
IL-6
MOCK
UV-JEV
JEV
Pg/mL
C
RANTES
Pg/ml
D
TNF-α
E
MCP-1
F
% Cell viability
G
Normalized JEV RNA levels
DMSO
Bromhexine
Clonidine
Flubendazole
Fluoxetine
LiCl
Memantine
Metformin
MTP
Rilmenidine
Sodium Valporate
H
IL-6
RANTES
TNF-α
IL-1α
DMSO
LPS
LPS+MTP


**Fig.S2**

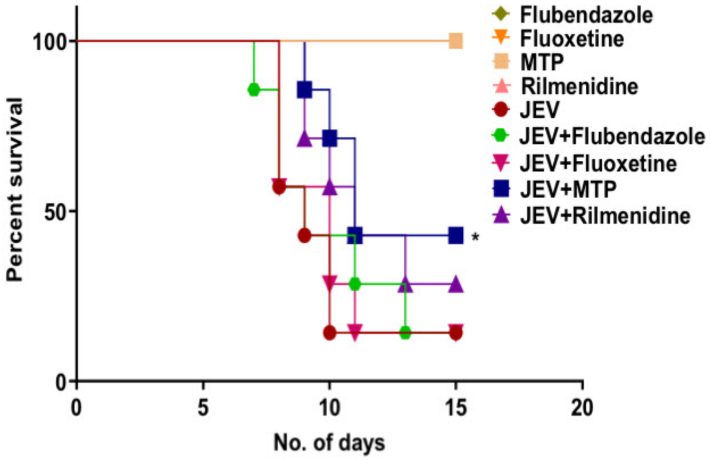
100
50
0
Percent survival
0
5
10
15
20
No. of days
*
Flubendazole
Fluoxetine
MTP
Rilmenidine
JEV
JEV+Flubendazole
JEV+Fluoxetine
JEV+MTP
JEV+Rilmenidine

**Fig.S3**

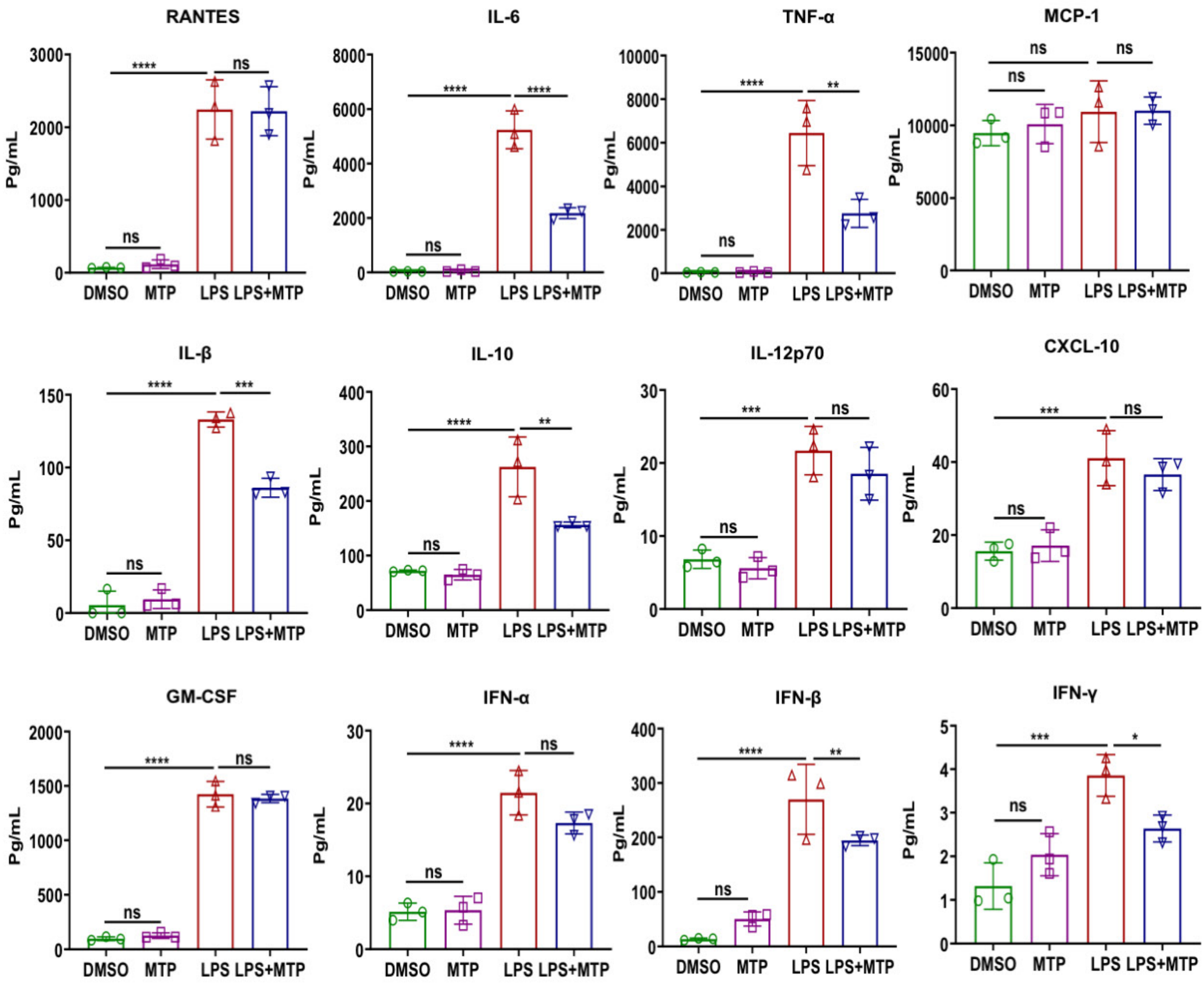
RANTES
IL-6
TNF-α
MCP-1
IL-β
IL-10
IL-12p70
CXCL-10
GM-CSF
IFN-α
IFN-β
IFN-γ
Pg/mL
DMSO
MTP
LPS
LPS+MTP
ns
****
***
**
*

**Fig.S4**

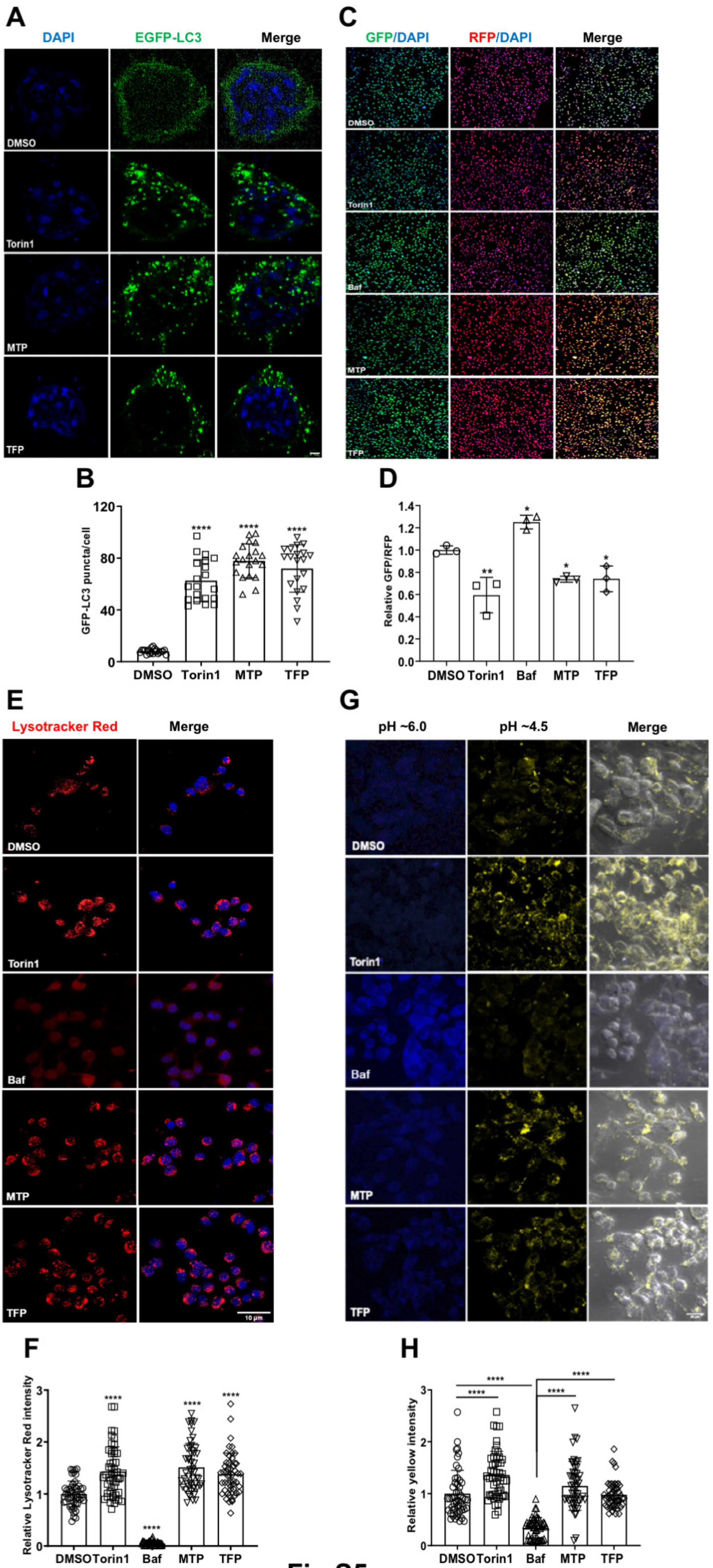
A
DAPI
EGFP-LC3
Merge
DMSO
Torin1
MTP
TFP
C
GFP/DAPI
RFP/DAPI
Merge
DMSO
Torin1
Baf
MTP
TFP
B
GFP-LC3 puncta/cell
0
40
80
120
****
****
****
DMSO
Torin1
MTP
TFP
D
Relative GFP/RFP
0.0
0.2
0.4
0.6
0.8
1.0
1.2
1.4
**
*
*
*
DMSO
Torin1
Baf
MTP
TFP
E
Lysotracker Red
Merge
DMSO
Torin1
Baf
MTP
TFP
10 μm
G
pH ~6.0
pH ~4.5
Merge
DMSO
Torin1
Baf
MTP
TFP
F
Relative Lysotracker Red intensity
0
1
2
3
****
****
****
****
DMSO
Torin1
Baf
MTP
TFP
H
Relative yellow intensity
0
1
2
3
****
****
****
****
DMSO
Torin1
Baf
MTP
TFP

Fig.S5

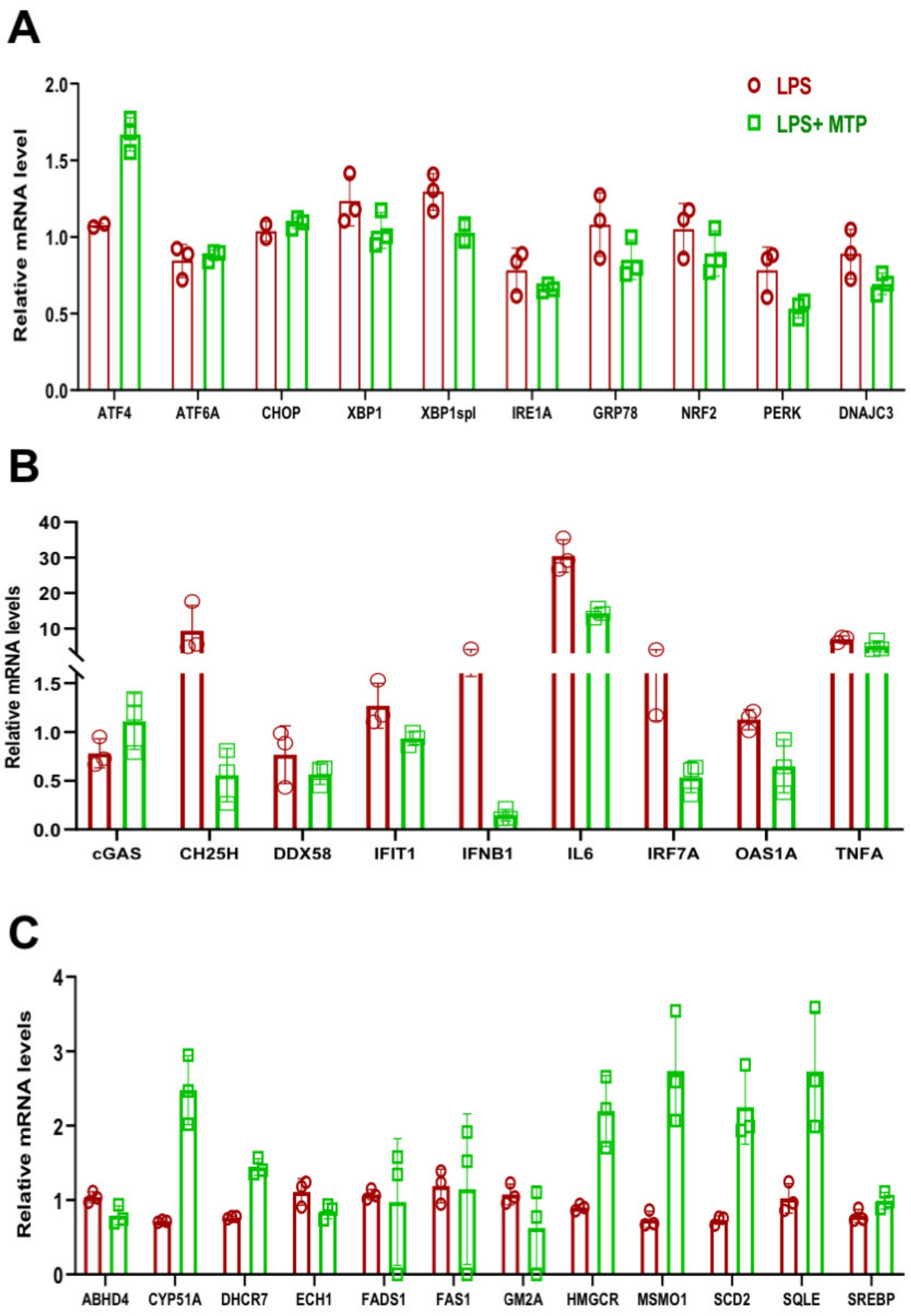

A
LPS
LPS+ MTP
Relative mRNA level
2.0
1.5
1.0
0.5
0.0
ATF4
ATF6A
CHOP
XBP1
XBP1spl
IRE1A
GRP78
NRF2
PERK
DNAJC3
B
Relative mRNA levels
40
30
20
10
1.5
1.0
0.5
0.0
cGAS
CH25H
DDX58
IFIT1
IFNB1
IL6
IRF7A
OAS1A
TNFA
C
Relative mRNA levels
4
3
2
1
0
ABHD4
CYP51A
DHCR7
ECH1
FADS1
FAS1
GM2A
HMGCR
MSMO1
SCD2
SQLE
SREBP


**Fig.S6**

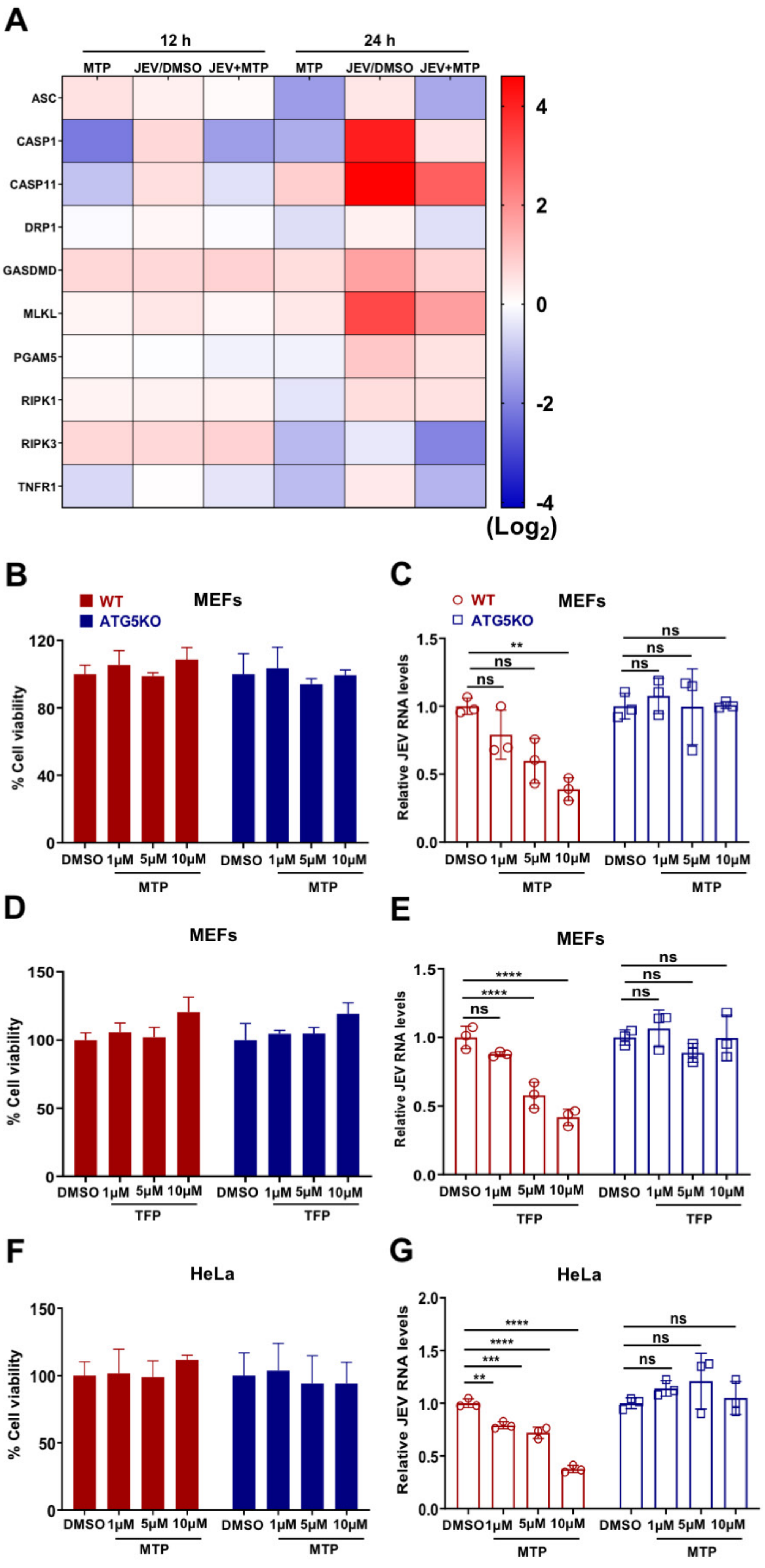
A
12 h
24 h
MTP
JEV/DMSO
JEV+MTP
ASC
CASP1
CASP11
DRP1
GASDMD
MLKL
PGAM5
RIPK1
RIPK3
TNFR1
4
2
0
-2
-4
(Log2)
B
WT
ATG5KO
MEFs
% Cell viability
DMSO 1μM 5μM 10μM
MTP
C
Relative JEV RNA levels
ns
**
D
TFP
E
****
F
HeLa
G
***

Fig.S7

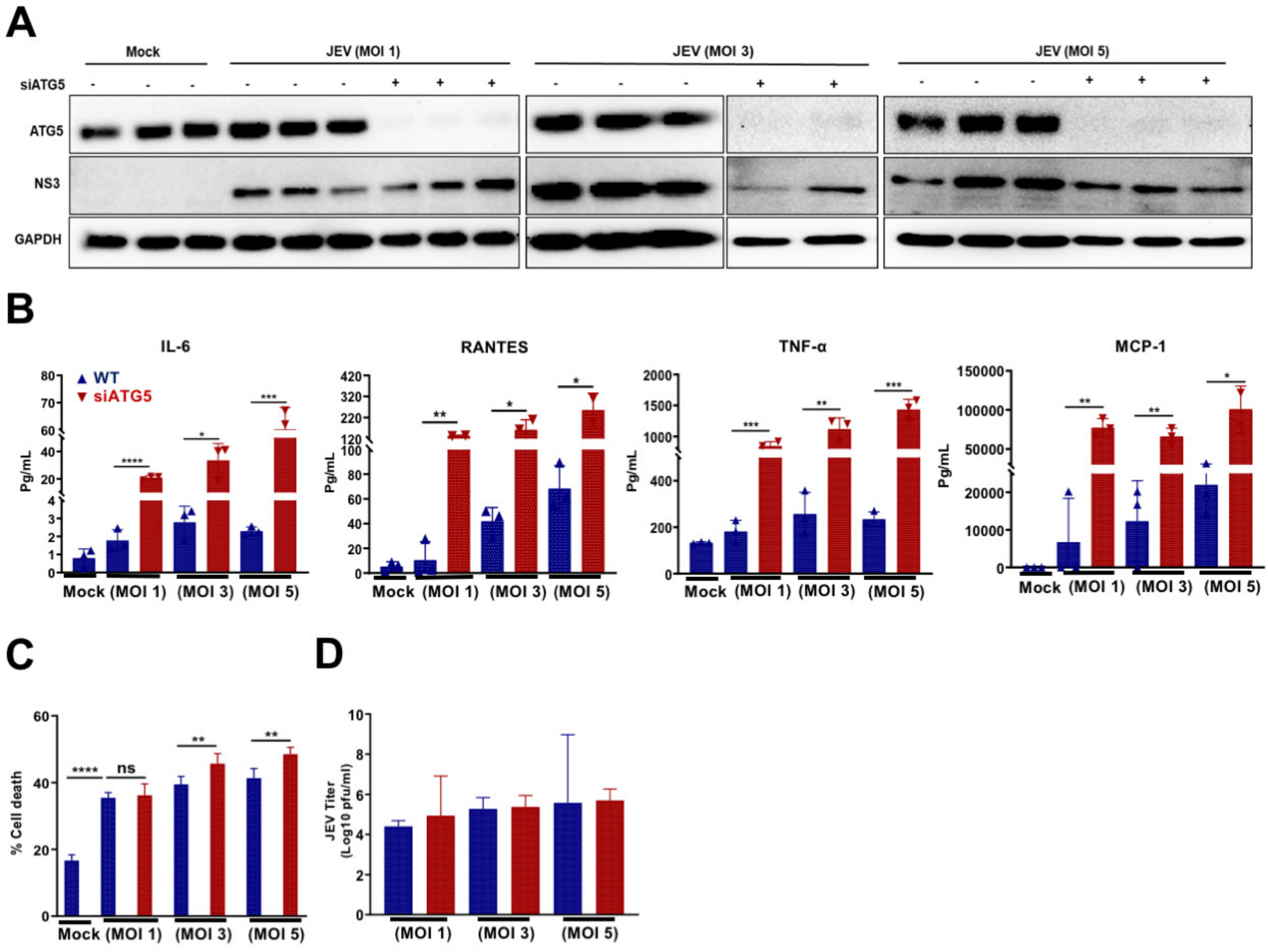
A
Mock
JEV (MOI 1)
JEV (MOI 3)
JEV (MOI 5)
siATG5
ATG5
NS3
GAPDH
B
IL-6
RANTES
TNF-α
MCP-1
WT
siATG5
Pg/mL
Mock
(MOI 1)
(MOI 3)
(MOI 5)
C
% Cell death
ns
D
JEV Titer
(Log10 pfu/ml)

Fig.S8

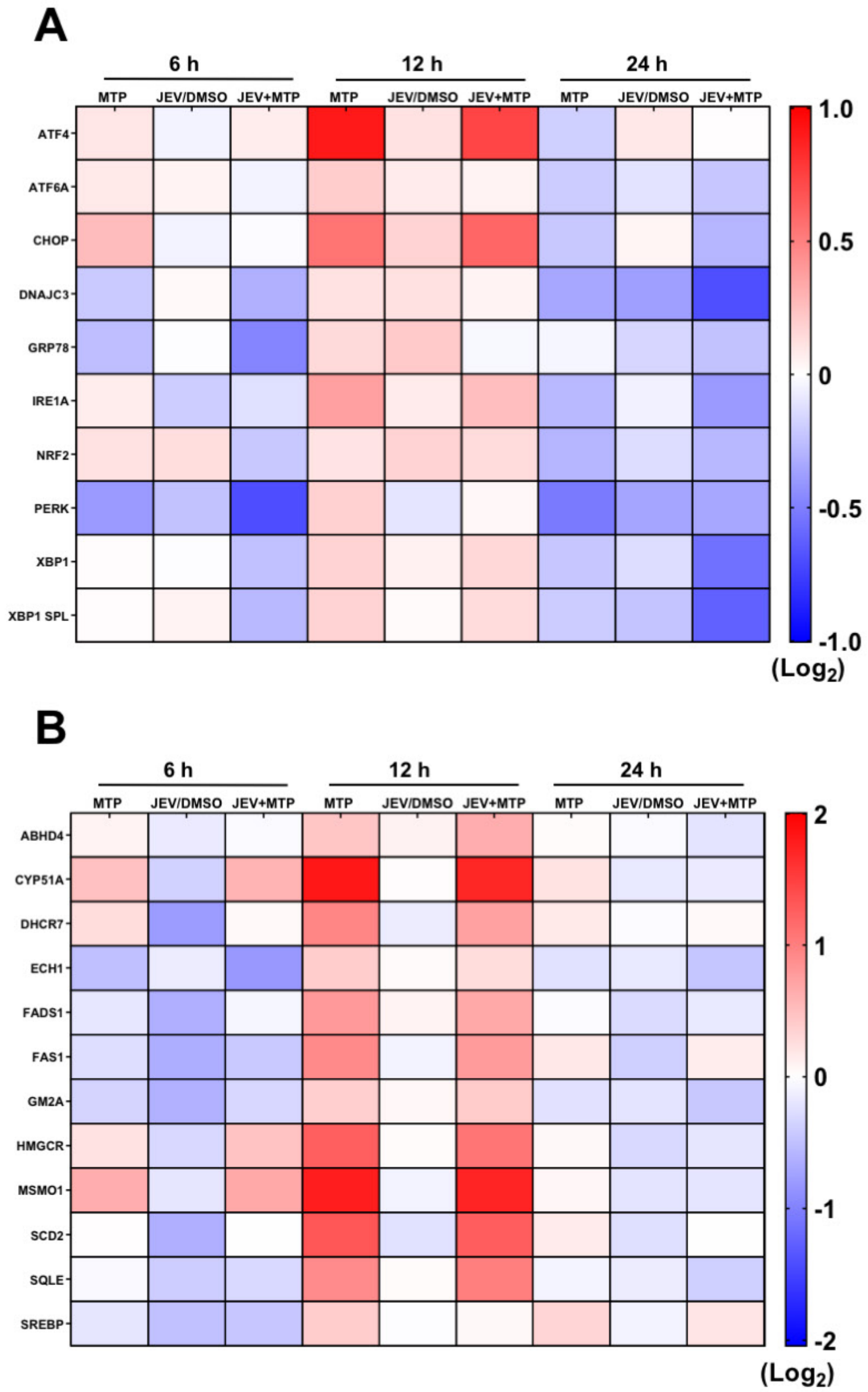
A
6 h
12 h
24 h
MTP
JEV/DMSO
JEV+MTP
ATF4
ATF6A
CHOP
DNAJC3
GRP78
IRE1A
NRF2
PERK
XBP1
XBP1 SPL
1.0
0.5
0
-0.5
-1.0
(Log2)
B
ABHD4
CYP51A
DHCR7
ECH1
FADS1
FAS1
GM2A
HMGCR
MSMO1
SCD2
SQLE
SREBP
2
1
0
-1
-2
(Log2)

**Fig.S9**

**Table S1: A panel of FDA-approved drugs used in this study**

| S.No. | FDA-approved Drug | References (PMID) |
|---|---|---|
| 1 | ABT-737 (BH3 mimetics) | 21460857 |
| 2 | Amiodarone hydrochloride | 18391949; 18024584 |
| 3 | Bromhexine | 26503418 |
| 4 | Carbamazepine | 18391949; 25535254 |
| 5 | Clomiphene citrate | 26503418 |
| 6 | Clonidine hydrochloride | 18391949 |
| 7 | Desipramine | 26503418; 24991837 |
| 8 | Diperodon | 26503418 |
| 9 | Diphenidol | 26503418 |
| 10 | Flubendazole | 26503418 |
| 11 | Fluoxetine | 26503418; 23364696 |
| 12 | Fluspirilene | 18024584 |
| 13 | Fluvoxamine | 26503418 |
| 14 | Hexachlorophene | 26503418 |
| 15 | Lithium chloride | 16186256 |
| 16 | L-NAME | 21726807 |
| 17 | Loperamide hydrochloride | 18391949 |
| 18 | Maprotiline | 26503418 |
| 19 | Melengestrol | 26503418 |
| 20 | Metformin | 21258367 |
| 21 | Memantine | 26503418 |
| 22 | Methotrimeprazine | 26503418 |
| 23 | Minoxidil | 18391949 |
| 24 | Nicardipine | 18024584 |
| 25 | Niguldipine | 18024584 |

| | | |
|---|---|---|
| 26 | Nimodipine | 18391949 |
| 27 | Nitrendipine | 18391949 |
| 28 | Nilvadipine | 26503418 |
| 29 | Nortriptyline | 26503418 |
| 30 | Oxiconazole | 26503418 |
| 31 | Pimozide | 18024584 |
| 32 | Proadifen | 26503418 |
| 33 | Resveratrol | 22465415 |
| 34 | Rilmenidine | 20190273; 18391949 |
| 35 | Sertraline | 26503418 |
| 36 | Sodium Valproate | 18391949; 25535254 |
| 37 | Statins-Pitavastatin | 24433351 |
| 38 | Ticlopidine | 26503418 |
| 39 | Trifluoperazine | 18024584 |
| 40 | Vitamin D | 21454634; 22589721 |
| 41 | Verapamil hydrochloride | 18391949 |
| 42 | Zotepine | 26503418 |

### TableS2: Chemical reagents used in this study

| S. No. | Name of Reagent | Catalogue No. |
|---|---|---|
| 1. | ABT-737 (BH3 mimetic) | Sigma (197333-10MG) |
| 2. | Agarose Type VII | Sigma (A4018) |
| 3. | Amiodarone hydrochloride | Sigma (PHR1164-1G) |
| 4. | Bafilomycin A1 | Sigma (B1793) |
| 5. | BCA assay kit | G-Biosciences (786-570) |
| 6. | Bromhexine | Sigma (PHR1831-200MG) |
| 7. | Carbamazepine | Sigma (PHR1067-1G) |
| 8. | CellTiter-Glo® CellTiter-Glo® Luminescent Cell Viability Assay | Promega (G7572) |
| 9. | Clomiphene citrate | Abcam (ab141183) |
| 10. | Clonidine hydrochloride | Sigma (C7897-100MG) |
| 11. | CM-H2DCFDA (General Oxidative Stress Indicator) | Invitrogen (C6827) |
| 12. | CyQUANT™ LDH Cytotoxicity Assay | Invitrogen (C20301) |
| 13. | Deoxyribonuclease I (DNase I) | SRL (61824) |
| 14. | Desipramine | Sigma (D3900-1G) |
| 15. | Dimethyl fumarate (DMF) | Sigma (242926-25G) |
| 16. | Dimethyl sulfoxide (DMSO) | Sigma (276855-250ML) |
| 17. | Diperodon hydrochloride | Sigma (D8536-5G) |
| 18. | Diphenidol hydrochloride | Sigma (SML2169-100MG) |
| 19. | EDTA | Sigma (E9884-500G) |
| 20. | Evans blue | Sigma (E2129-10G) |
| 21. | Flubendazole | Abcam (ab143260) |
| 22. | Fluoxetine hydrochloride | Sigma (F132-10MG) |
| 23. | Fluspirilene | Sigma (F100-10MG) |

| | | |
|---|---|---|
| 24. | Fluvoxamine maleate | Abcam (ab141082) |
| 25. | Fura-2, AM | Invitrogen (F1225) |
| 26. | Glucose | Sigma (G8270-100G) |
| 27. | Haloperidol | Sigma (H1512-5G) |
| 28. | HBSS | Gibco (14175095) |
| 29. | HEPES | Sigma (H3375-25G) |
| 30. | Hexachlorophene pestanal | Sigma (45526-250MG) |
| 31. | ImProm-II™ Reverse Transcription System | Promega (A3800) |
| 32. | Lithium chloride | Abcam (ab120853) |
| 33. | L-NAME | Abcam (ab120136) |
| 34. | Loperamide hydrochloride | Cayman Chemical (14875) |
| 35. | LPS | Sigma (L2630) |
| 36. | LysoSensor™ Yellow/Blue DND-160 | Invitrogen (L7545) |
| 37. | LysoTracker™ Red DND-99 | Invitrogen (L7528) |
| 38. | Maprotiline (hydrochloride) | Cayman Chemical (15892) |
| 39. | Melengestrol acetate | Sigma (33998-100MG-R) |
| 40. | Memantine hydrochloride | Sigma (M9292-25MG) |
| 41. | Metformin | Abcam (ab120847) |
| 42. | Methotrimeprazine | MedChem Express (HY-B1693) |
| 43. | Minoxidil | Sigma (M4145-25MG) |
| 44. | MTT 3-(4,5-Dimethylthiazol-2-yl)-2,5-Diphenyltetrazolium Bromide | VWR life science, (0793-1G) |
| 45. | N-Acetyl-L-cysteine (NAC) | Sigma (A7250-10G) |
| 46. | Nicardipine hydrochloride | Sigma (N7510-1G) |
| 47. | Niguldipine (hydrochloride) | Cayman Chemical (19534) |
| 48. | Nilvadipine | Sigma (SML0945-10MG) |
| 49. | Nimodipine | Sigma (N149-100MG) |

| | | |
|---|---|---|
| 50. | Nitrendipine | Sigma (N144-25MG) |
| 51. | Nortriptyline (hydrochloride) | Cayman Chemical (15904) |
| 52. | Oxiconazole | Sigma (SML1474-10MG) |
| 53. | Phenylmethylsulfonyl fluoride (PMSF) | Sigma (329-98-6) |
| 54. | Pimozide | Abcam (ab142135) |
| 55. | Poly (ethylene glycol) (PEG 400) | Sigma (202398-500G) |
| 56. | Premix Ex Taq™ (Probe qPCR) | Takara (RR390A) |
| 57. | Proadifen | Sigma (P1061-100MG) |
| 58. | ProLong™ Gold Antifade Mountant with DAPI | Invitrogen (P36935) |
| 59. | Protease inhibitor cocktail (PI) | Sigma (P8340) |
| 60. | Puromycin | InvivoGen (ant-pr-1) |
| 61. | PVDF membrane | Merck Millipore (IPVH00010) |
| 62. | Random hexamer | Sigma (H0268) |
| 63. | RBC lysis buffer | GCC Biotech (19114B1076) |
| 64. | Resveratrol | Abcam (ab120726) |
| 65. | Rilmenidine hemifumarate salt | Sigma (R134-5MG) |
| 66. | SDS | Sigma (L3771-500G) |
| 67. | Sodium pyruvate | Himedia (TCL015) |
| 68. | Sodium Valproate | Abcam (ab120745) |
| 69. | Statins-Pitavastatin | Cayman Chemical (15414) |
| 70. | SYBR® Premix Ex Taq™ | Takara (RR420A) |
| 71. | Thapsigargin | Sigma (T9033) |
| 72. | Ticlopidine | Sigma (T6654-1G) |
| 73. | Torin1 | Tocris Bioscience (4247) |
| 74. | Trifluoperazine Dihydrochloride | Cayman Chemical (15068) |
| 75. | Triton™ X-100 | Sigma (T9284-500ML) |

| | | |
|---|---|---|
| 76. | Trizol reagent (RNAiso Plus) | Takara (9109) |
| 77. | Tween 20 | G-Biosciences (RC1227) |
| 78. | Verapamil hydrochloride | Abcam (ab120140) |
| 79. | Vitamin D | Cayman Chemical (11791) |
| 80. | Zotepine | Sigma (Z0877-10MG) |
| | **Media & other additives** | **Catalogue No.** |
| 81. | 2XMEM | Himedia (AL178A-500ML) |
| 82. | B-27 | Gibco (17504044) |
| 83. | DMEM | Himedia (AL007A-500ML) |
| 84. | FBS | Himedia, (RM10432-500ML) |
| 85. | Geneticin™ Selective Antibiotic (G418 Sulfate) | Gibico (10131035) |
| 86. | L-15 | Himedia (AL011S-500ML) |
| 87. | L929 conditioned media | Culture supernatant of L929 fibroblast cells (After 6-8 days starvation of L929 cells) |
| 88. | L-Glutamine | Himedia (TCL012) |
| 89. | MEM | Hyclone (SH3024401-500ML) |
| 90. | Neurobasal | Gibco (21103049) |
| 91. | Penicillin-Streptomycin | Himedia (A007-100ML) |
| 92. | RPMI-1640 | Himedia (AL028-500ML) |
| 93. | Trypsin - EDTA Solution | Himedia (TCL007) |
| | **Antibodies** | **Catalogue No.** |
| 94. | Alexa fluor™ 488 chicken anti-mouse IgG (H+L) | Invitrogen (A-21200) |
| 95. | Alexa fluor™ 568 donkey anti-mouse IgG (H+L) | Invitrogen (A-10037) |
| 96. | ATG5 | CST (12994S) |
| 97. | GAPDH | GeneTex (GTX100118) |

| | | |
|---|---|---|
| 98. | JEV-Envelope | Abcam (ab41671) |
| 99. | JEV-NS1 | Abcam (ab41651) |
| 100. | LC3B | Abcam (ab51520) |
| 101. | mTOR | CST (2983S) |
| 102. | p-4E-BP1 (Thr37/46) | CST (2855S) |
| 103. | p70S6Kinase | CST (9202S) |
| 104. | Peroxidase AffiniPure Donkey Anti-Mouse IgG (H+L) | Jackson ImmunoResearch (715-035-150) |
| 105. | Peroxidase AffiniPure Donkey Anti-Rabbit IgG (H+L) | Jackson ImmunoResearch (711-035-152) |
| 106. | p-mTOR (Ser2448) | CST (5536S) |
| 107. | p-p70S6Kinase (Thr389) | CST (97596S) |
| 108. | SQSTM1/ p62 | Abcam (ab56416) |
| | **Plasmids** | **Catalogue No.** |
| 109. | gag/pol | Addgene (14887) |
| 110. | pCI-VSVG | Addgene (1733) |
| 111. | pEGFP-LC3 | Addgene (21073) |
| 112. | pEGFP-N1-TFEB | Addgene (38119) |
| 113. | pMRX-IP-GFP-LC3-RFP-LC3ΔG | Addgene (84572) |
| | **siRNA/Transfection reagents** | **Catalogue No.** |
| 114. | DharmaFECT 2 | Dharmacon (T-2002-02) |
| 115. | Lipofectamine™ RNAimax | Invitrogen (13778030) |
| 116. | Lipofectamine™ 2000 | Invitrogen (11668019) |
| 117. | ON-TARGETplus Non-targeting (NT) | Dharmacon (D-001810-10-20) |
| 118. | ON-TARGETplus Mouse siAtg5 | Dharmacon (L-064838-00-0005) |
| 119. | ON-TARGETplus Mouse siAtg7 | Dharmacon (L-049953-00-0005) |
| | **Cytokines** | **Catalogue No.** |

| | | |
|---|---|---|
| 120. | Mouse IL-1α Flex Set | BD Bioscience (560157) |
| 121. | Mouse IL-1β Flex Set | BD Bioscience (560232) |
| 122. | Mouse IL-6 Flex Set | BD Bioscience (558301) |
| 123. | LEGENDPLEX MU Anti-Virus Response Panel (13-plex) | Biolegend (740622) |
| 124. | Mouse MCP-1 Flex Set | BD Bioscience (558342) |
| 125. | Mouse/Rat Soluble Protein Master Buffer Kit | BD Bioscience (558266) |
| 126. | Mouse RANTES Flex Set | BD Bioscience (558345) |
| 127. | Mouse TNF Flex Set | BD Bioscience (558299) |

### TableS3: Primers used in this study

| S.No. | Gene name | Forward (5'-3') sequence | Reverse (5'-3') sequence |
|---|---|---|---|
| 1. | ABHD4 | GGCACAGTTTGGGAGGATTCC | ACTAGGGTCAGTTGGTCGTAG |
| 2. | AKT2 | ATGAACGACGTAGCCATTGTG | TTGTAGCCAATAAAGGTGCCAT |
| 3. | AMPK/PRKAA1 | GTCAAAGCCGACCCAATGATA | CGTACACGCAAATAATAGGGGTT |
| 4. | ASC | CTTGTCAGGGGATGAACTCAAAA | GCCATACGACTCCAGATAGTAGC |
| 5. | ATF4 | CTCTTGACCACGTTGGATGAC | CAACTTCACTGCCTAGCTCTAAA |
| 6. | ATF6 A | CGGTCCACAGACTCGTGTTC | GCTGTCGCCATATAAGGAAAGG |
| 7. | ATG12 | TCCCCGGAACGAGGAACTC | TTCGCTCCACAGCCCATTTC |
| 8. | ATG13 | CCAGGCTCGACTTGGAGAAAA | AGATTTCCACACACATAGATCGC |
| 9. | ATG14 | GAGGGCCTTTACGTGGCTG | AATAGACGAAATCACCGCTCTG |
| 10. | ATG16L1 | CAGAGCAGCTACTAAGCGACT | AAAAGGGGAGATTCGGACAGA |
| 11. | ATG3 | ACACGGTGAAGGGAAAGGC | TGGTGGACTAAGTGATCTCCAG |
| 12. | ATG4A | GCTGGTATGGATTCTGGGGAA | TGGGTTGTTCTTTTTGTCTCTCC |
| 13. | ATG4B | TATGATACTCTCCGGTTTGCTGA | GTTCCCCCAATAGCTGGAAAG |
| 14. | ATG5 | TGTGCTTCGAGATGTGTGGTT | GTCAAATAGCTGACTCTTGGCAA |
| 15. | ATG7 | GTTCGCCCCCTTTAATAGTGC | TGAACTCCAACGTCAAGCGG |
| 16. | ATG9A | CAGTTTGACACTGAATACCAGCG | AATGTGGTGCCAAGGTGATTT |
| 17. | BCL2 | ATGCCTTTGTGGAACTATATGGC | GGTATGCACCCAGAGTGATGC |
| 18. | BECN1 | ATGGAGGGGTCTAAGGCGTC | TCCTCTCCTGAGTTAGCCTCT |
| 19. | CASP1 | ACAAGGCACGGGACCTATG | TCCCAGTCAGTCCTGGAAATG |
| 20. | CASP11 | ACAAACACCCTGACAAACCAC | CACTGCGTTCAGCATTGTTAAA |
| 21. | CGAS | GAGGCGCGGAAAGTCGTAA | TTGTCCGGTTCCTTCCTGGA |
| 22. | CH25H | TGCTACAACGGTTCGGAGC | AGAAGCCCACGTAAGTGATGAT |
| 23. | CHOP/DDIT3 | ACCTTCACTACTCTTGACCCTG | GATGTGCGTGTGACCTCTGT |
| 24. | CYP51A1 | GACAGGAGGCAACTTGCTTTC | GTGGACTTTTCGCTCCAGC |
| 25. | DDX58 | AAGAGCCAGAGTGTCAGAATCT | AGCTCCAGTTGGTAATTTCTTGG |
| 26. | DHCR7 | AGGCTGGATCTCAAGGACAAT | GCCAGACTAGCATGGCCTG |
| 27. | DNAJC3 | GGCGCTGAGTGTGGAGTAAAT | GCGTGAAACTGTGATAAGGCG |
| 28. | DRP1 | CAGGAATTGTTACGGTTCCCTAA | CCTGAATTAACTTGTCCCGTGA |
| 29. | ECH1 | GCTACCGCGATGACAGTTTC | TCAGAGATCGAAGGCTGATGTT |
| 30. | EIF2AK2/PKR | ATGCACGGAGTAGCCATTACG | TGACAATCCACCTTGTTTTCGT |
| 31. | FADS1 | AGCACATGCCATACAACCATC | TTTCCGCTGAACCACAAAATAGA |
| 32. | FASN F | GGAGGTGGTGATAGCCGGTAT | TGGGTAATCCATAGAGCCCAG |
| 33. | GADD34/PPP1R15A | GCCTGCAAGGGGCTGATAAG | TTTGTATCCCGGAGCTATGGA |
| 34. | GAPDH | CGTCCCGTAGACAAAATGGT | TTGATGGCAACAATCTCCAC |
| 35. | GM2A | CGCCTTTCCCAACTTGGTG | TGACGACTACATCTCCAGGAAC |
| 36. | GRP78/HSPA5 | GCATCACGCCGTCGTATGT | ATTCCAAGTGCGTCCGATGAG |
| 37. | GSDMD | CCATCGGCCTTTGAGAAAGTG | ACACATGAATAACGGGGTTTCC |
| 38. | HGAPDH | TGCACCACCAACTGCTTACG | GGCATGGACTGTGGTCATGAG |
| 39. | HMGCR | AGCTTGCCCGAATTGTATGTG | TCTGTTGTGAACCATGTGACTTC |
| 40. | IFIT1 | CTGAGATGTCACTTCACATGGAA | GTGCATCCCCAATGGGTTCT |

| | | | |
|---|---|---|---|
| 41. | IFNB1 | CAGGTAGTAGGCGACACTGT | TCAATTGCCACAGGAGCTTC |
| 42. | IFN-α | ATGGCTAGRCTC TGTGCTTTCCT | AGGGCTCTCCAGAYTTCTGCTCTG |
| 43. | IFN-β | AAGAGTTACACTGCCTTTGCCATC | CACTGTCTGCTGGTGGAGTTCATC |
| 44. | IFN-γ | GGCCATCAGCAACATAAGCGT | TGGGTTGTTGACCTCAAACTTGGC |
| 45. | IL-6 | CTGCAAGAGACTTCCATCCAG | AGTGGTATAGACAGGTCTGTTGG |
| 46. | IRE1 ALPHA/*ERN1* | ACACCGACCACCGTATCTCA | CTCAGGATAATGGTAGCCATGTC |
| 47. | IRF7 | GAGACTGGCTATTGGGGGAG | GACCGAAATGCTTCCAGGG |
| 48. | JEV | AGAGCACCAAGGGAATGAAATAGT<br>Taqman probe: CCACGCCACTCGACCCATAGACTG (5′ end FAM, 3′end TAMRA). | AATAAGTTGTAGTTGGGCACTCTG |
| 49. | LAMP1 | CAGCACTCTTTGAGGTGAAAAAC | ACGATCTGAGAACCATTCGCA |
| 50. | LAMP2 | TGTATTTGGCTAATGGCTCAGC | TATGGGCACAAGGAAGTTGTC |
| 51. | LC3A | GACCGCTGTAAGGAGGTGC | CTTGACCAACTCGCTCATGTTA |
| 52. | LC3B | TTATAGAGCGATACAAGGGGGAG | CGCCGTCTGATTATCTTGATGAG |
| 53. | MCP-1/CCL2 | CAAGAAGGAATGGGTCCAGA | GCTGAAGACCTTAGGGCAGA |
| 54. | MDA5 | AGATCAACACCTGTGGTAACACC | CTCTAGGGCCTCCACGAACA |
| 55. | MLKL | AATTGTACTCTGGGAAATTGCCA | TCTCCAAGATTCCGTCCACAG |
| 56. | MSMO1 | AAACAAAAGTGTTGGCGTGTTC | AAGCATTCTTAAAGGGCTCCTG |
| 57. | mTOR | ACCGGCACACATTTGAAGAAG | CTCGTTGAGGATCAGCAAGG |
| 58. | NRF2/NFE212 | TAGATGACCATGAGTCGCTTGC | GCCAAACTTGCTCCATGTCC |
| 59. | OAS1 | GCCTGATCCCAGAATCTATGC | GAGCAACTCTAGGGCGTACTG |
| 60. | PERK/E*IF2AK3* | GCACTTTAGATGGACGAATCGC | TGCTGAGGCTAGATGAAACCA |
| 61. | PGAM5 | ATCTGGAGAAGACGAGTTGACA | CCTGTTCCCGACCTAATGGT |
| 62. | PI3KCA | CCACGACCATCTTCGGGTG | ACGGAGGCATTCTAAAGTCACTA |
| 63. | PRKAA2 | CAGGCCATAAAGTGGCAGTTA | AAAAGTCTGTCGGAGTGCTGA |
| 64. | RANTES/CCL5 | GCTGCTTTGCCTACCTCTCC | TCGAGTGACAAACACGACTGC |
| 65. | RIG-I | ACAGATCCGAGACACTAAAGGG | AACAGCGCCTCTGATGGAAAG |
| 66. | RIPK1 | GAAGACAGACCTAGACAGCGG | CCAGTAGCTTCACCACTCGAC |
| 67. | RIPK3 | TCTGTCAAGTTATGGCCTACTGG | GGAACACGACTCCGAACCC |
| 68. | SCD2 | GCATTTGGGAGCCTTGTACG | AGCCGTGCCTTGTATGTTCTG |
| 69. | SQLE | ATAAGAAATGCGGGGATGTCAC | ATATCCGAGAAGGCAGCGAAC |
| 70. | SREBP2 | GCAGCAACGGGACCATTCT | CCCCATGACTAAGTCCTTCAACT |
| 71. | TLR-3 | GTGAGATACAACGTAGCTGACTG | TCCTGCATCCAAGATAGCAAGT |
| 72. | TNFR1sf1a | CCGGGAGAAGAGGGATAGCTT | TCGGACAGTCACTCACCAAGT |
| 73. | TNF-α | CCCTCACACTCAGATCATCTTCT | GCTACGACGTGGGCTACAG |
| 74. | ULK1 | AAGTTCGAGTTCTCTCGCAAG | CGATGTTTTCGTGCTTTAGTTCC |
| 75. | VPS34/PIK3C3 | CCTGGACATCAACGTGCAG | TGTCTCTTGGTATAGCCCAGAAA |
| 76. | XBP-1 | AGCAGCAAGTGGTGGATTTG | GAGTTTTCTCCCGTAAAAGCTGA |
| 77. | XBP-1 SPLICED | GACAGAGAGTCAAACTAACGTGG | GTCCAGCAGGCAAGAAGGT |
